# Acclimation of carbon metabolism to a changing environment across a leaf rosette of *Arabidopsis thaliana*

**DOI:** 10.1101/2025.04.29.651223

**Authors:** Vladimir Brodsky, Anian Kerscher, Michaela Urban, Thomas Nägele

**Affiliations:** LMU München, Faculty of Biology, Plant Evolutionary Cell Biology, Großhaderner Str. 2-4, 82152 Planegg, Germany

**Keywords:** *Arabidopsis* thaliana, cold acclimation, leaf development, photosynthesis, carbon metabolism

## Abstract

- Plants need to efficiently and quickly stabilize photosynthesis and carbon metabolism under changing environmental conditions to prevent irreversible tissue damage
- Differently developed leaves of *Arabidopsis thaliana* rosettes are typically homogenized for metabolic analyses which might result in significant over- or underestimation of metabolite dynamics.
- Here, photosynthesis and carbon metabolism were analysed in mature and immature leaves of rosettes of natural Arabidopsis accessions originating from southern and northern Europe. Ambient growth condition at 22 °C was compared to a combined low temperature/elevated light treatment.
- Gradients of F_v_/F_m_ and CO_2_ assimilation rates across leaf rosettes hinted towards tissue-specific acclimation capacities of photosynthesis and carbon metabolism.
- Dynamics of carbohydrates and carboxylic acids were integrated with photosynthetic parameters in a quantitative carbon balance model. Model simulations suggested that mature leaf tissue stabilizes acclimation of carbon metabolism in immature leaf tissue and/or other sink tissue.
- In conclusion, acclimation capacities of photosynthesis to low temperature and elevated light significantly differ within one leaf rosette of Arabidopsis. This directly affects carbon metabolism which shapes the metabolic acclimation capacity of the whole plant.
- It is suggested to consider such tissue-specific effects which might otherwise be hidden behind a high variance of experimental data.

## Introduction

Plant metabolism is highly plastic and allows for efficient buffering of environmental dynamics (Shaar-Moshe et al., 2019). Central to environmentally induced metabolic adjustments is the balance of primary and secondary photosynthetic reactions, i.e., balancing of photosynthetic light energy absorption and biochemical carbon fixation within the Calvin-Benson-Bassham cycle (CBBc). Carbohydrates are the direct products of photosynthetic CO_2_ fixation, and regulation of their metabolism plays a central role in plant acclimation to environmental changes (Rolland et al., 2006; Ruan et al., 2010). In photosynthetically active leaf tissue, triose phosphates are synthesised in the CBBc and provide carbon equivalents for biosynthesis of transitory starch and sucrose. In a changing light and/or temperature regime, the balance of carbon fluxes towards storage compounds, e.g., starch, and soluble sugars was found to be of central importance for the acclimation capacity. For example, enzymatic deficiency in the starch degradation pathway was reported to result in impaired freezing tolerance which was accompanied by a significantly affected metabolism and accumulation response of soluble sugars (Yano et al., 2005). Similarly, starch deficiency resulted in a delay of hexose accumulation during cold exposure which suggested a role of starch degradation in augmentation of soluble sugars during early cold response (Sicher, 2011). Particularly, during the early period of cold exposure, such augmentation might be essential to sustain energy metabolism until adjustments in transcription, translation and metabolic regulation have stabilised metabolism.

Sucrose biosynthesis, catalysed by sucrose phosphate synthase (SPS) enzymes, has been shown earlier to essentially contribute to photosynthetic cold acclimation (Strand et al., 2003; Nägele et al., 2012). Fructose 6-phosphate and UDP-glucose are substrates for the SPS reaction which yields sucrose 6-phosphate being immediately dephosphorylated by sucrose phosphate phosphatase (SPP). Sucrose might serve as a substrate for carbon export into sink organs or can be hydrolyzed by invertase enzymes which are located in different compartments, e.g., the cytosol and vacuole (Sturm, 1999; Roitsch and González, 2004; Nägele, 2022). Glucose and fructose, which are products of invertase reactions, are substrates for hexokinases which, in an ATP-dependent manner, phosphorylate both hexoses to produce glucose 6-phosphate (G6P) and fructose 6-phosphate (F6P), respectively (Granot et al., 2013). As G6P is substrate for UDP-glucose biosynthesis, these reactions constitute a potential cycle of sucrose cleavage and re-synthesis which has been discussed before to be involved in carbon partitioning and stabilization of carbohydrate metabolism against environmental perturbation (Geigenberger and Stitt, 1991; Fürtauer and Nägele, 2016).

Carbohydrates are well known to be essentially involved in metabolic and photosynthetic regulation in a changing environment, but physiological interpretation of carbohydrate dynamics is frequently challenged due to their multiple roles in metabolism, signaling and development (Moore et al., 2003; Ruan, 2014). For example, due to being a substrate for hydrolysis and carbon export from source to sink tissue, sucrose dynamics might basically be explained in different ways: either due to a changed balance between biosynthesis and hydrolysis or between biosynthesis and export rates. Further, sink-source relationships between plant tissues might change in the life cycle of a plant which simultaneously affects expression and activity patterns of enzymes and transporters involved in sugar metabolism (Koch, 2004; Durand et al., 2018). Additionally, cyclic interconversion of metabolites, such as sucrose, makes analysis of observed carbohydrate dynamics non-intuitive due to an increased number of explanatory models of metabolism. To support the analysis and interpretation, mathematical modelling of metabolite dynamics has been proven useful in previous studies (Stelling et al., 2002; Shi and Schwender, 2016; Töpfer et al., 2020). Frequently, ordinary differential equation (ODE) systems are applied for kinetic modelling and simulation of metabolite dynamics over time. In such a model, ODEs are used to describe metabolite dynamics by the sum of in- and output fluxes which might comprise biosynthesis, interconversion or transport processes. In a kinetic model, these processes are typically described by enzyme kinetic terms which contain experimentally determined enzyme parameters and variables, e.g., values of K_M_, K_i_ and v_max_. Finally, model simulation, i.e., numerical integration of the ODE system, results in metabolite concentrations which can be evaluated experimentally.

In the present study, we have combined experimental analysis of photosynthesis and carbohydrate metabolism with kinetic modelling to reveal effects of cold and elevated light on metabolic regulation across old (mature) and young (immature) leaf tissue in natural accessions of *Arabidopsis thaliana*. The accessions originated from northern and southern Europe reflecting a broad geographical range of European *Arabidopsis* populations. Based on an observed significant gradient of maximum quantum yield of Photosystems II between old and young leaves under low temperature and elevated light, it was hypothesized that this was also reflected in carbon metabolism. Differential dynamics of carbohydrates and carboxylic acids further supported a tissue-specific photosynthetic and metabolic acclimation process.

## Materials and methods

### Plant material and growth conditions

Natural accessions of *Arabidopsis thaliana*, Ct-1 (geographical origin: Catania, Italy; NASC ID: N6674), Fei-0 (geographical origin: St. Maria d. Feiria, Portugal; NASC ID: N22645), Oy-0 (geographical origin: Oystese, Norway; NASC ID: N6824) and Rsch-4 (geographical origin: Rschew/Starize, Russia; NASC ID: N6850), were grown on a 1:1 mixture of GS90 soil and vermiculite in a climate chamber under controlled short-day conditions (8 h/16 h light/dark; 100 μmol m^−2^ s^−1^; 22°C/16°C; 50-60% relative humidity). After eight weeks, plants were sampled at the end of night (eon), and at the end of day (eod). For treatment, plants were transferred to a cold room for low temperature and elevated light treatment (LT/EL; 8 h/16 h light/ dark; 250 μmol m^−2^ s^−1^; 4-6°C/4°C, 40-50% relative humidity). After 7 days at LT/EL, plants were sampled at eon and eod. Young and old leaves were immediately quenched separately in liquid nitrogen and stored for up to three months at -60°C until further use. A schematic overview of sampled young/old tissue is provided in the supplements (**Supplementary Figure S1**).

### Net photosynthesis and chlorophyll fluorescence measurements

Net rates of CO_2_ assimilation (NPS) were recorded under growth light intensities, i.e., 100 or 250 μmol m^−2^ s^−1^, at midday, with a WALZ GFS-3000FL system (Heinz Walz GmbH; www.walz.com).

Parameters of chlorophyll fluorescence were obtained through Imaging-PAM MAXI version at ambient temperature (Heinz Walz GmbH; www.walz.com). Maximum quantum yield of PSII (F_v_/F_m_) was determined after 15 minutes of dark adaptation of the rosette and subsequently supplying a saturating light pulse.

### Quantification of starch and soluble carbohydrates

Transitory starch and soluble carbohydrate amounts were determined as described before (Kitashova et al., 2023). Fresh leaves mixed with sea sand were homogenized in 80% ethanol and incubated at 80°C for 30 minutes. The supernatant, containing the soluble sugars, was separated from the starch-containing pellet and transferred to a separate tube for further analysis. Pellets were hydrolyzed in 0.5 N NaOH at 95°C for 60 min and acidified with 1 M CH_3_COOH. The suspension was then subjected to amyloglucosidase digestion. Resulting glucose moieties were quantified photometrically at 540 nm in a coupled glucose-oxidase/peroxidase/dianisidine assay.

Ethanol of previously obtained soluble sugar fractions was evaporated and the dried residue solubilized in water. Sucrose was quantified by incubating the samples in 30% KOH at 95°C for 10 minutes before adding 0.14% (w/v) anthrone in 14.6 M H_2_SO_4_ and incubating at 40°C for 30 min. Complexes were photometrically detected and quantified with a calibration curve at 620 nm.

Glucose amounts were obtained in a coupled enzymatic assay of hexokinase and glucose-6-phosphate dehydrogenase (G6PDH), while fructose amounts were quantified via a coupled hexokinase/phosphoglucose isomerase/G6PDH assay. In both cases, NADP^+^ was reduced to yield NADPH+H^+^ which was quantified at 340nm.

### Quantification of enzyme activities

Enzyme activities under optimal temperature and substrate saturation (v_max_) were obtained for sucrose phosphate synthase (SPS), glucokinase (GLCK), fructokinase (FRCK), and invertase (INV) as described previously (Kitashova et al., 2023).

Leaf tissue was homogenized with sea sand in 50 mM HEPES pH 7.5, 10 mM MgCl_2_, 1 mM EDTA, 2.5 mM DTT, 10% (v/v) glycerine and 0.1% (v/v) Triton X-100. SPS enzymes were extracted on ice for 20 minutes. Following centrifugation at 4°C with 20,000 g, the supernatant was incubated for 30 min at 25°C with 50 mM HEPES pH 7.5, 15 mM MgCl_2_, 2.5 mM DTT, 35 mM UDP-glucose, 35 mM F6P and 140 mM G6P. The activity assay was stopped by boiling samples with 30% KOH and sucrose amounts were determined with an anthrone assay as described above.

For glucokinase (GLCK) and fructokinase (FRCK) enzyme activity assays, homogenized leaf tissue was extracted in 50 mM Tris pH 8.0, 0.5 mM MgCl_2_, 1 mM EDTA, 1 mM DTT and 1% (v/v) Triton X-100. Samples were centrifuged at 4°C with 20,000 g and the supernatant was combined with a buffer containing 100 mM HEPES pH 7.5, 10 mM MgCl_2_, 2 mM ATP, 1 mM NADP^+^, 0.5 U G6PDH. Activities were determined from changes of NADPH+H^+^ over time, resulting from addition of 5 mM Glucose or Fructose, for respective measurements of v_max:GLCK_ and v_max:FRCK_, at 30°C and 340 nm.

Cytosolic (nINV) and vacuolar (aINV) invertase activity was determined in 50 mM HEPES–KOH pH 7.5, 5 mM MgCl_2_, 2 mM EDTA, 1 mM phenylmethylsulfonylfluoride, 1 mM DTT, 10% (v/v) glycerine and 0.1% (v/v) Triton X-100 on ice. The suspension was centrifuged at 4°C with 20,000 g incubated in (I), an nInv-specific reaction buffer with a pH of 7.5, containing 20 mM HEPES–KOH and 100 mM sucrose, or (II), in an aInv-specific reaction buffer at ph 4.7, containing 20 mM sodium acetate and 100 mM sucrose. After sufficiently long incubation at 30°C, the reactions were terminated at 95°C. Similar to the starch assay, cleaved glucose was then determined photometrically at 540 nm by a coupled glucose oxidase/peroxidase/dianisidine assay.

### Quantification of carboxylic acids

Absolute amounts of citrate, fumarate and malate were determined using photometric assay kits (Sigma-Aldrich, MAK057, MAK060 and MAK067). Carboxylic acids were extracted in hot water, i.e., at 95°C for 15 min. After brief centrifugation, the supernatant was used for photometric measurements.

### Data evaluation and mathematical modelling

Data was statistically analysed in R Version 3.6.2 (www.r-project.org) (R Core Team 2019) and RStudio Version 1.2.5033 (www.rstudio.com) (RStudio Team 2019). Mathematical modelling was performed in MATLAB Version 24.2.0.2740171 (R2024b) (www.mathworks.com) with the IQM Tools Toolbox, developed by IntiQuan (www.intiquan.com) and the curve fitting toolbox.

For parameter estimation, a particle swarm pattern search method for bound constrained global optimization was utilized (Vaz and Vicente, 2007). A cost function was minimized which quantified squared errors between simulations and experimental data.

The model described a simplified approximation of the central leaf carbohydrate metabolism of *Arabidopsis thaliana.* It was assumed that carbon equivalents can be exchanged between young and old leaves (**Figure 1**). To simulate the metabolic environment, a set of ordinary differential equations was used to describe daily dynamics of metabolite concentrations. Experimentally determined enzyme activities were used to simulate Michaelis-Menten kinetics with regulatory parameters K_i_ and K_m_ derived from a previous study (Nägele et al., 2010). All models are provided in the supplements (**Supplementary Data F1**).

**Figure 1.**
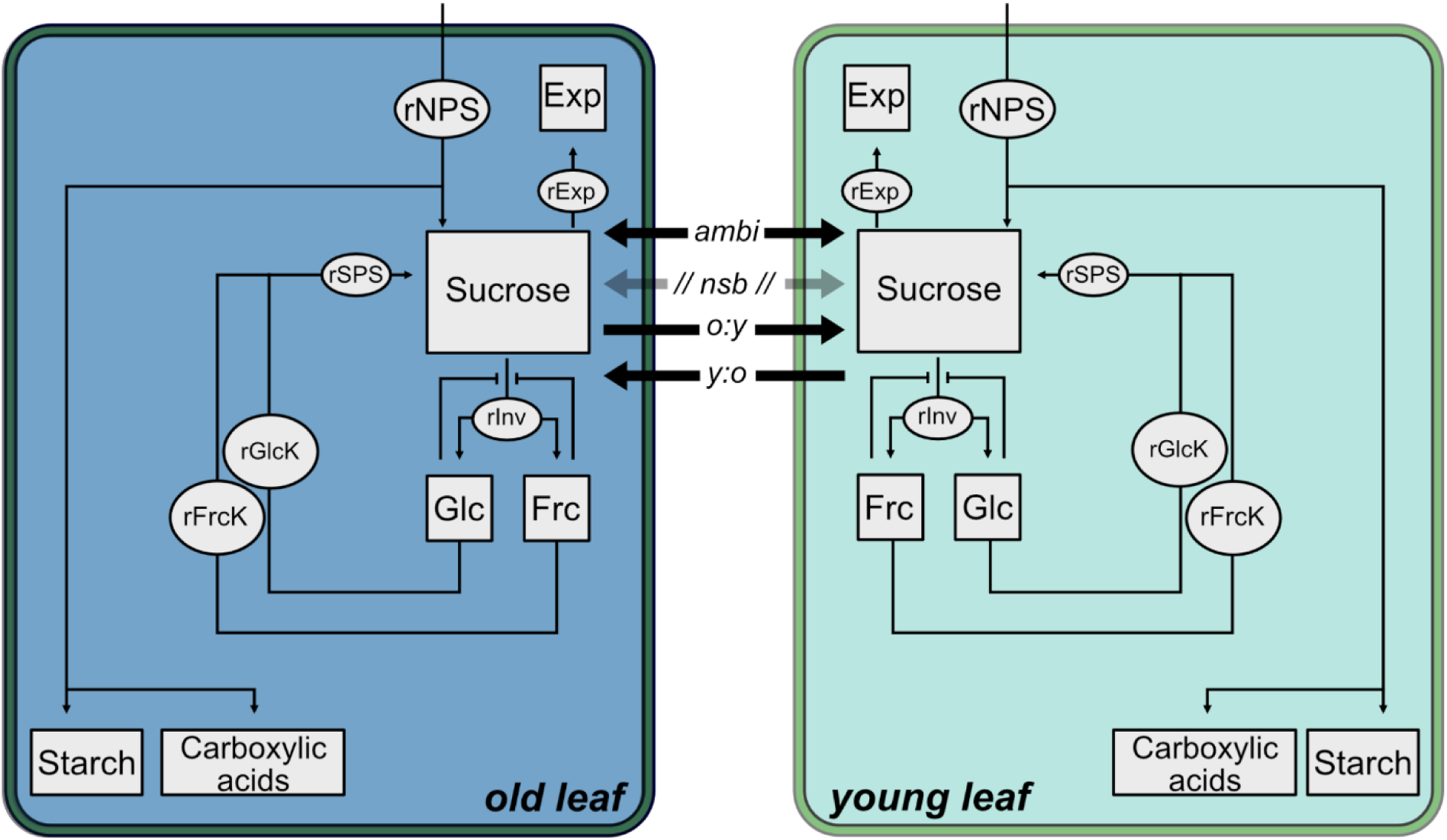
Model of putative metabolic interactions between carbon metabolism in young and old leaf tissue. Exp: unspecific sucrose efflux; Glc: glucose; Frc: fructose; *rNPS*, rate of net photosynthesis; rSPS, rate of sucrose phosphate synthase; rInv, rate of invertase; rGlck, rate of glucokinase; rFrck, rate of fructokinase; *ambi*: ambidirectional sucrose transport; *nsb*: no sucrose transport between tissues; *o:y*: sucrose transport from old to young tissue; *y:o*: sucrose transport from young to old tissue. Arrows with blunt ends indicate feedback inhibition by products, i.e., feedback inhibition of invertases by hexoses.

In simulations of metabolism under low temperature, enzymatic rates were adjusted to the temperature regime by multiplication with the correction factor T_Arrh_, which was selected from the interval I_temp_ = [0.25; 0.4]. This factor was described previously as the reduction of maximal enzymatic activity resulting from thermodynamic constraints acting on the enzyme at 4°C. It was obtained by logarithmic conversion of the Arrhenius equation and providing the desired experimental temperature (Kitashova et al., 2023).

The system input function was described by NPS carbon assimilation rates. From these rates, transitory starch and the dynamics of summed carboxylic acids (i.e., citrate, fumarate, malate) were subtracted. This metric was quantified by adding the median of diurnal starch accumulation to the mean of carboxylic acid accumulation and calculating the hourly accumulation rate assuming a linear rate across the light phase, i.e., 8 hours. This value was subtracted from the mean of the carbon assimilation rate and stoichiometrically adjusted to represent the rate of carbon-input in C_6_ units. Calculated uptake rates were compared to the maximal activity of SPS enzymes. Due to the finding that the sucrose input was verified to be lower than the SPS v_max_ under all conditions, these model assumptions were maintained.

For carbohydrate and enzyme data, parameter bounds were set to the 25th and 75th quartiles derived from the experimental data. In the case of invertase activity, v_max_ of acidic and neutral invertases were summarized and parametric bounds were constrained to the range from lower to upper standard deviation from the mean.

Maximal bounds for a putative export of sucrose were set empirically for each accession x condition. Models that did not contain a sucrose bridging point between old and young leaves (nsb) had their upper bound of sucrose export (*k_exp*) adjusted in integer steps, until a solution was found that could explain experimentally determined data. This *k_exp* was then fixed for the other models of the same accession x condition and the bounds for export of sucrose into the other leaf were kept as the same as the ones for *k_exp*.

## Results

### Photosynthesis is affected by leaf development

The analysis of maximum quantum yield of Photosystem II (F_v_/F_m_) revealed a significant developmental effect of leaf tissue during cold acclimation across all tested natural accessions Ct-1, Fei-0, Oy-0 and Rsch-4 (**Figure 2**). Generally, F_v_/F_m_ values did not differ significantly between leaf tissues under ambient temperature. Yet, upon exposure to LT/EL, F_v_/F_m_ values of old leaves were observed to be significantly lower than in young tissue (ANOVA, p < 0.05).

**Figure 2.**
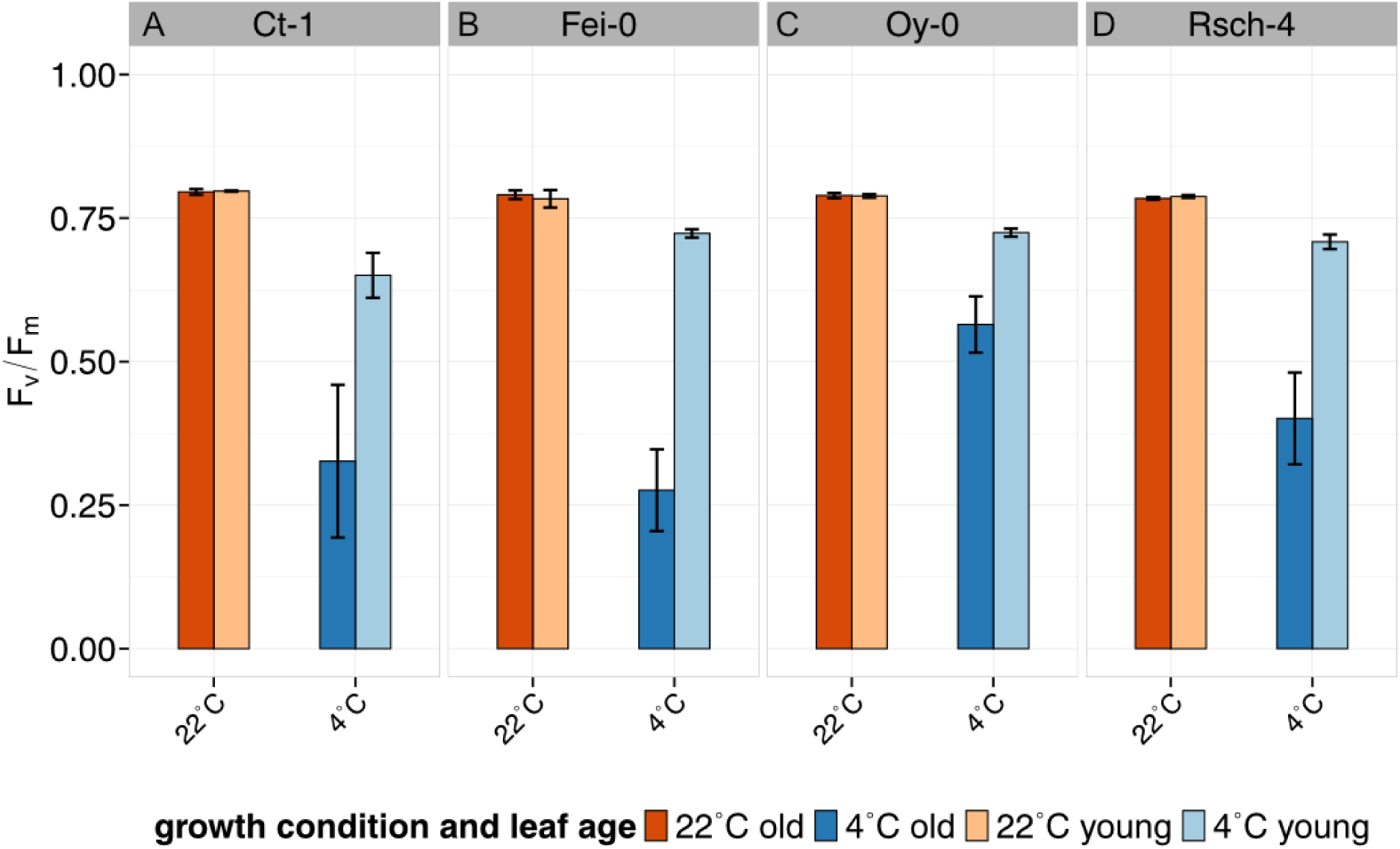
Maximum quantum yield of Photosystem II (F_v_/F_m_) at low temperature and elevated light. **(A)** Ct-1, **(B)** Fei-0, **(C)** Oy-0, **(D)** Rsch-4. Bars represent means ± SD, n = 3. Orange: ambient temperature conditions, blue: 7d of LT/EL. Light shade: young leaves, dark shade: old leaves.

To evaluate if these differences were related to carbon assimilation, rates of net photosynthesis (NPS) were quantified. All NPS rates were quantified at 22 °C applying the respective growth PAR intensities, i.e., 100 µmol photons m^−2^ s^−1^ for control plants and 250 μmol photons m^−2^ s^−1^ for LT/EL plants. At 22 °C, young leaves had a lower NPS than old leaves, except for Fei-0 where no difference was detected between both tissues (**Figure 3**; ANOVA, p < 0.05). Consistently across all accessions, under LT/EL, mean rates of NPS in young leaves were found to be higher than in old leaves while this effect was only found to be significant in Ct-1 (**Figure 3 A**; ANOVA, p < 0.05).

**Figure 3.**
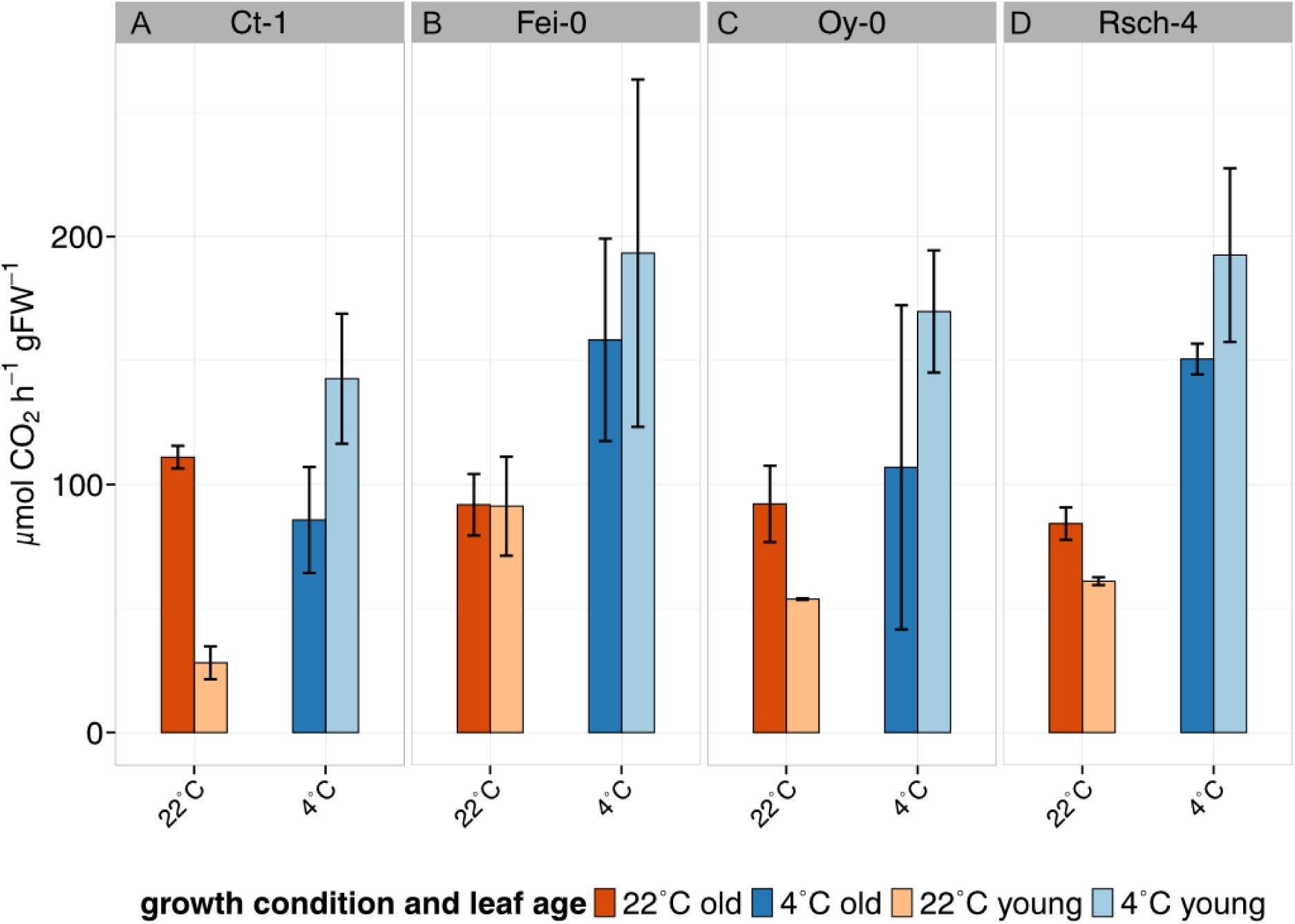
Rates of net carbon assimilation across all tested accessions, tissues and growth conditions. **(A)** Ct-1, **(B)** Fei-0, **(C)** Oy-0, **(D)** Rsch-4. Rates were determined at 22°C and growth PAR intensity, i.e., at 100 μmol photons m^−2^ s^−1^ for control plants and at 250 μmol photons m^−2^ s^−1^ for LT/EL plants. Bars represent means ± SD, n = 3. Orange: ambient temperature conditions, blue: 7d of LT/EL. Dark shade: old leaves; Light shade: young leaves.

Comparing both F_v_/F_m_ and NPS in a principal component analysis (PCA) revealed a separation of both growth regimes along principal component (PC) 1, capturing 65.1% of the total variance (**Figure 4**). The separation was more pronounced for young than for old tissue. Both tissue types were separated along PC2 which captured ∼36% of total variance. The (approximately) perpendicular orientation of both loadings, i.e., F_v_/F_m_ and NPS, revealed that they were not correlated. This showed that NPS was not (linearly) predictable from F_v_/F_m_, and *vice versa*.

**Figure 4.**
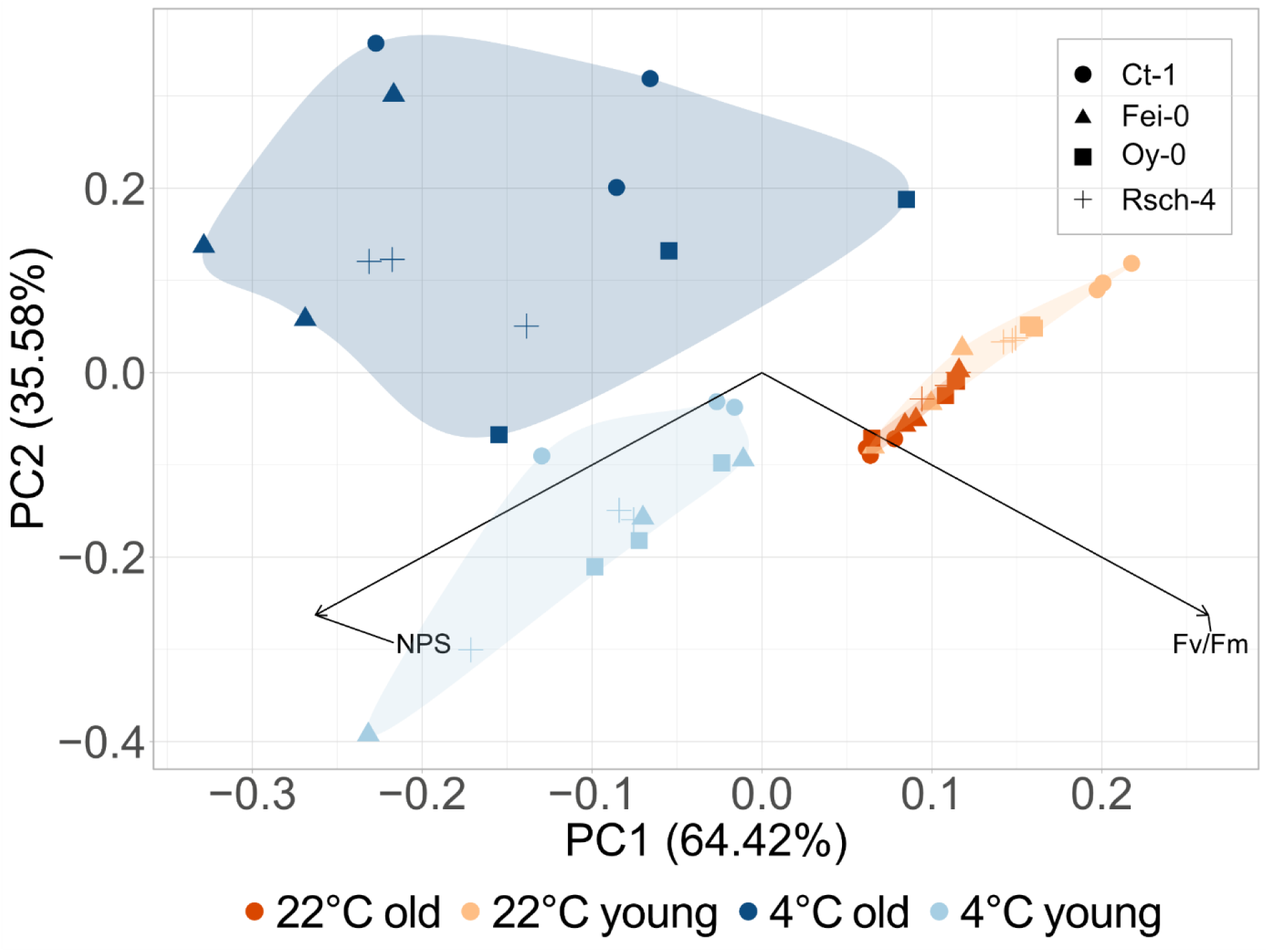
Principal component analysis of Fv/Fm and rates of NPS across accessions, growth conditions and leaf tissues. Parameters were determined at 22°C and growth PAR intensity, i.e., at 100 μmol photons m^−2^ s^−1^ for control plants and at 250 μmol photons m^−2^ s^−1^ for LT/EL plants. Orange: ambient temperature conditions, blue: 7d of LT/EL. Light shade: young leaves, dark shade: old leaves. Shapes indicate accessions: Ct-1: circles; Fei-0: triangles; Oy-0: squares; Rsch-4: crosses.

### Leaf development affects dynamics of central pools of carbon metabolism

Sucrose and transitory starch are central products of photosynthetic CO_2_ fixation and represent an essential branching point for carbon allocation within leaf metabolism. Consistently, across accessions, tissue types and conditions, both carbohydrates were observed to increase during the light phase (**Figure 5**). In all accessions, starch accumulated significantly during the day at 22 °C in both old and young leaf tissue (**Figure 5 A-D**). At LT/EL, the accessions differed in their starch accumulation pattern: Ct-1 showed a significant starch accumulation only in young tissue (**Figure 5 A**), while no diurnal accumulation was observed in Fei-0 anymore (**Figure 5 B**). In Oy-0 and Rsch-4, both young and old tissue showed a significant starch accumulation (**Figure 5 C, D**).

**Figure 5.**
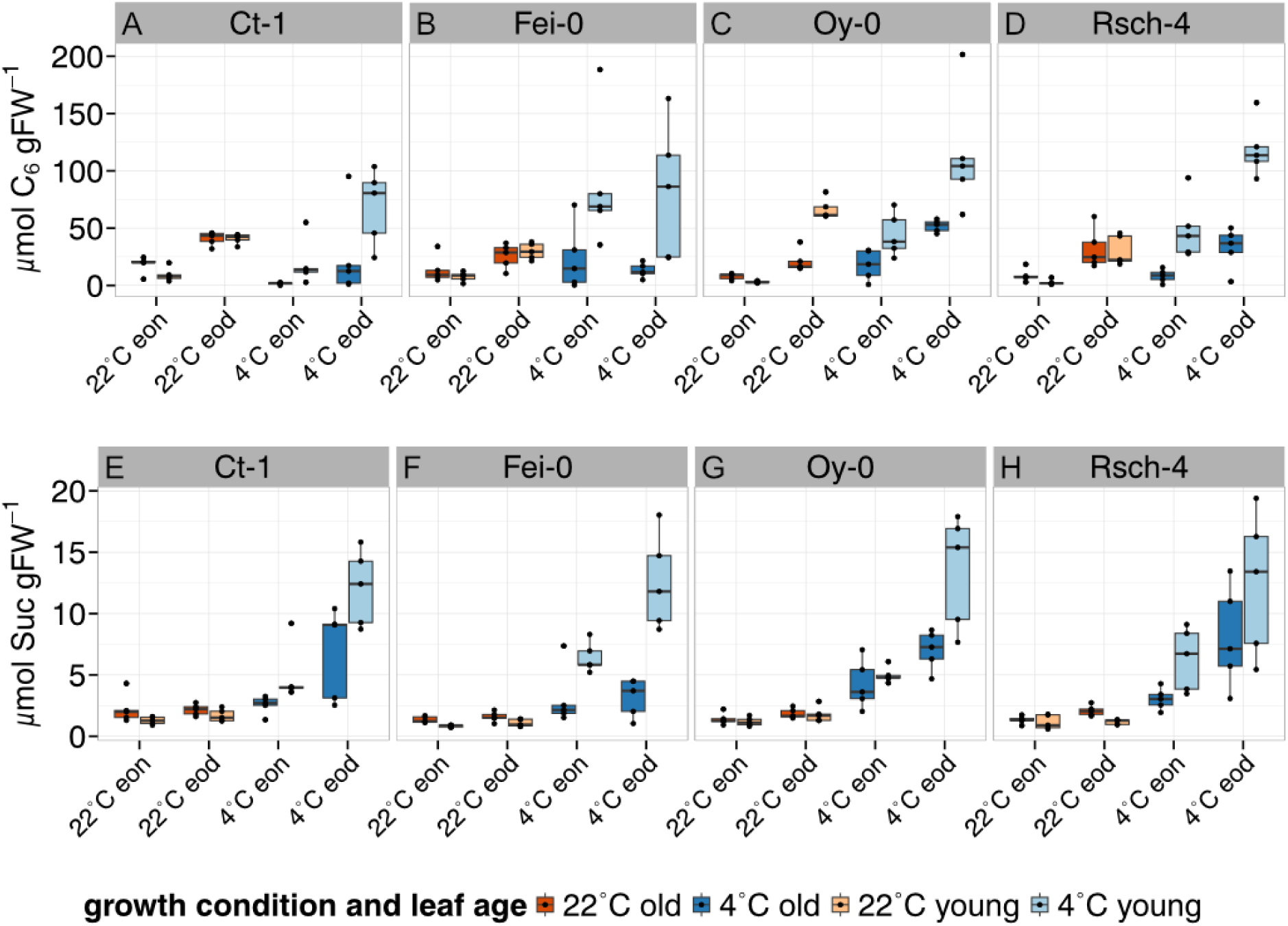
Diurnal dynamics of starch and sucrose amounts. **(A) – (D)** Diurnal dynamics of starch amounts, expressed as C_6_ moieties, in old and young tissue of natural accessions before (22C) and after (4C) LT/EL treatment. **(E) - (H)** Diurnal dynamics of sucrose amounts in old and young tissue of natural accessions before (22C) and after (4C) LT/EL treatment. Orange: ambient temperature conditions, blue: 7d of LT/EL. Light shade: young leaves, dark shade: old leaves. n = 5.

Under ambient conditions, sucrose amounts were found to be similar across accessions and leaf tissues (**Figure 5 E-H**). At LT/EL, sucrose amounts were observed to be elevated in all leaf tissues of all accessions. Further, diurnal sucrose accumulation was pronounced under LT/EL and resulted in highest levels in young leaf tissue.

Under ambient conditions, the amounts of free hexoses glucose and fructose showed slight dynamics during the day which was, however, different between accessions (**Figure 6**). While Ct-1 showed constant glucose amounts, Fei-0 and Oy-0 were found to increase their glucose amounts until end of the day (**Figure 6 A - C**). More consistently, under LT/EL, the amounts of both hexoses increased in all accessions. Under these conditions, glucose was found to accumulate during the light phase, particularly in young tissue, whereas fructose amounts remained constant (**Figure 6 E-H**).

**Figure 6.**
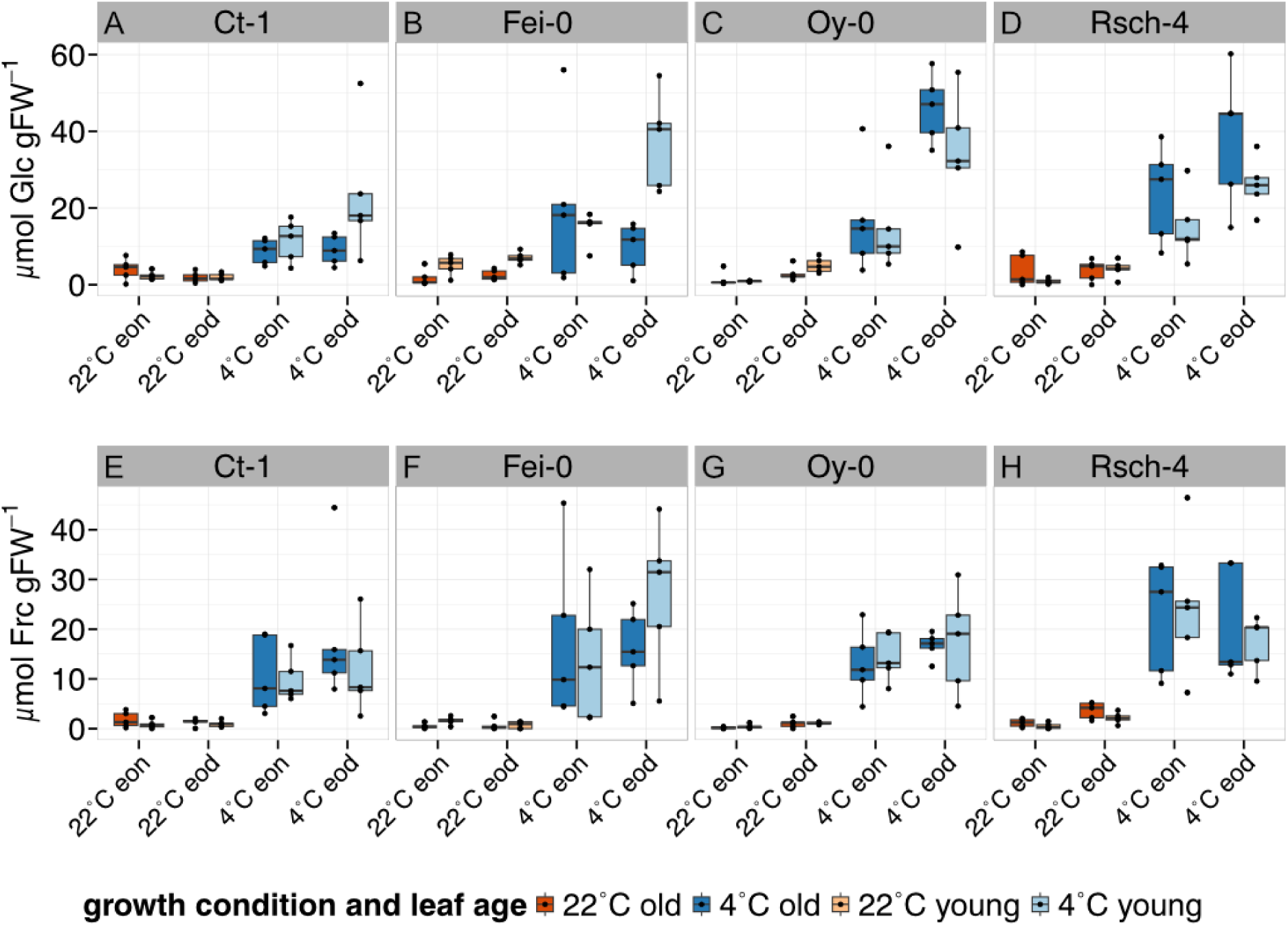
Dynamics of glucose and fructose amounts. **(A) - (D)** Diurnal dynamics of glucose in old and young tissue of natural accessions before (22C) and after (4C) LT/EL treatment. **(E) - (H)**: Diurnal dynamics of fructose in old and young tissue of natural accessions before (22C) and after (4C) LT/EL treatment. Orange: ambient temperature conditions, blue: 7d of LT/EL. Light shade: young leaves, dark shade: old leaves. n = 5.

To reveal whether exposure to LT/EL affected metabolism of carboxylic acids, amounts of citrate, fumarate and malate were quantified (**Figure 7, Supplementary Figures S2-S4**). At 22 °C, the amounts of carboxylic acid in old leaves were approximately twice as high as in young leaves. Exposure to LT/EL resulted in a similar total amount of carboxylic acids in young and old leaves. Only in Fei-0, amounts in young leaves under LT/EL were lower than in old leaves (**Figure 7 B**).

**Figure 7.**
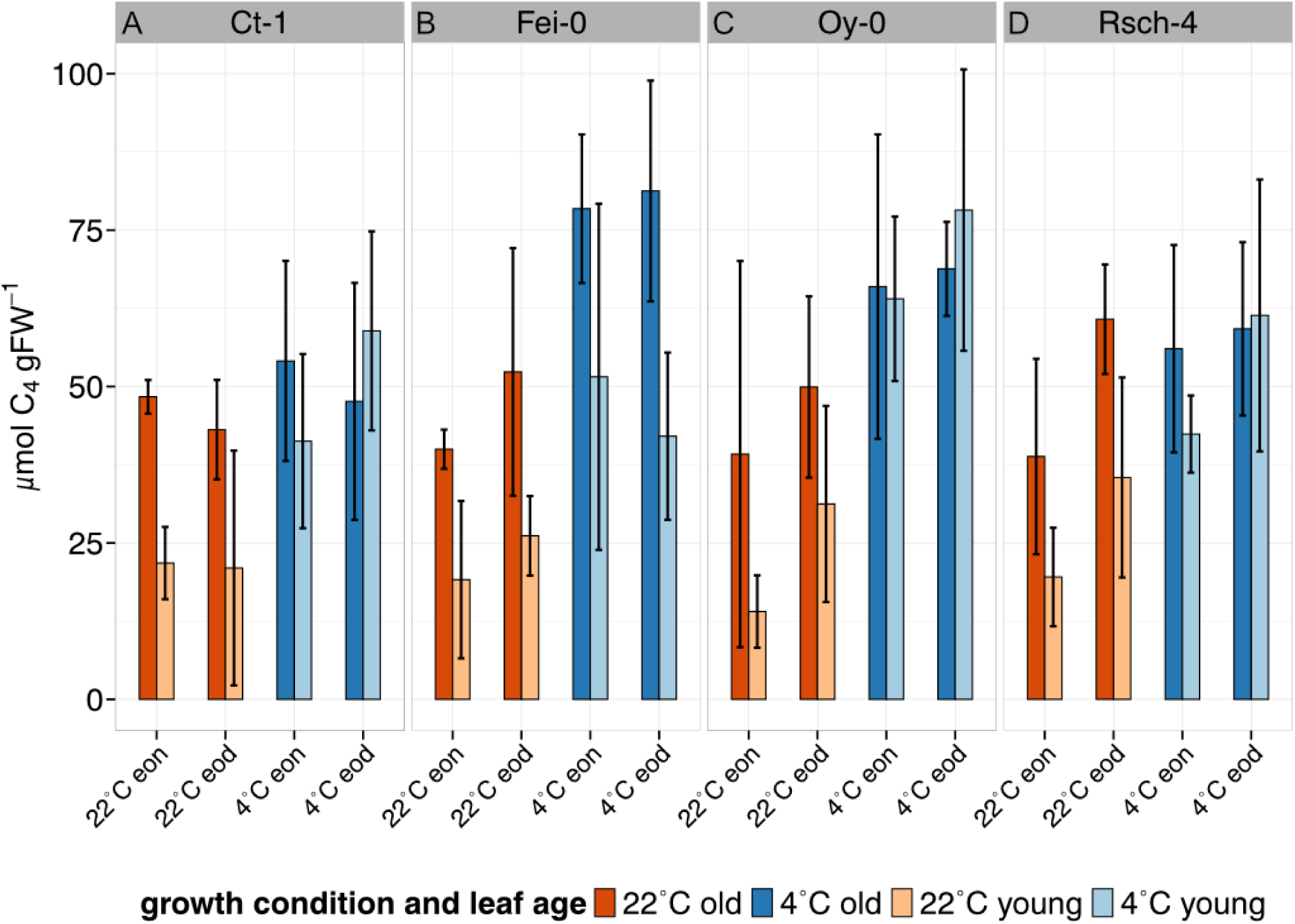
Diurnal dynamics of carboxylic acids. Summed quantities (in C4 units) of citrate, fumarate and malate for **(A)** Ct-1, **(B)** Fei-0, **(C)** Oy-0, **(D)** Rsch-4. Bars represent means ± SD, n = 3. Orange: ambient temperature conditions, blue: 7d of LT/EL. Light shade: young leaves, dark shade: old leaves. A summary of each carboxylic acid is provided in the supplements (**Supplementary Figures S2 – S4**).

### Carbon balance relates differently to metabolism in young and old leaf tissue

Net carbon fluxes during the light phase were estimated by balancing carbon uptake rates (NPS), carbon turnover (C(to)) and carbon export to other tissue and/or pathways (EXP). C(to) was calculated by stoichiometrically converting metabolite pools to C6 equivalents and assessing their (summed) accumulation rates per hour during the 8-hour light phase. EXP was then derived as the difference between NPS and C(to). A Pearson correlation analysis revealed that young and old tissue showed a different relation between metabolites and enzymes to these estimated carbon fluxes under environmental changes which suggested differential regulation of metabolism and photosynthesis (**Figure 8, Supplementary Figure S5**).

**Figure 8.**
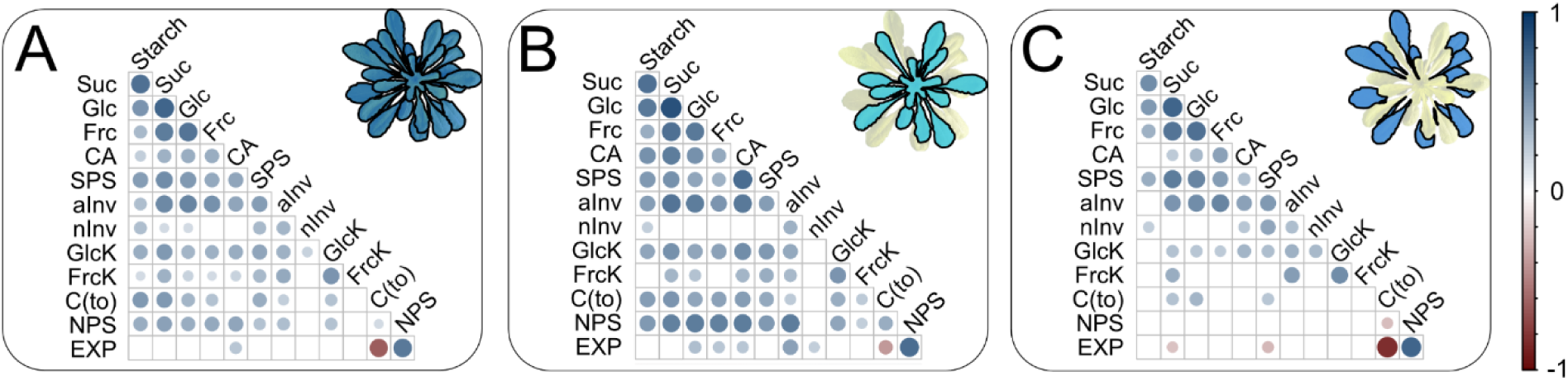
Pearson correlation of metabolites, enzyme activities and estimated carbon fluxes in young and old leaf tissue. **(A)** Pearson correlation across all tissue, time points and accessions. **(B)** Pearson correlation across young tissue, time points and accessions. (C) Pearson correlation across old tissue, time points and accessions. Colorbar indicates positive (blue) and negative (red) correlation coefficients. Blank fields: non-significant correlations (p ≥ 0.05). Suc: sucrose; Glc: glucose; Frc: fructose; CA: carboxylic acids (i.e., summed citrate, malate and fumarate); SPS: sucrose phosphate synthase; aInv: acidic invertase; nInv: neutral invertase; GlK: glucokinase; FrK: fructokinase; C(to): carbon turnover rates; NPS: net photosynthesis rates; EXP: calculated export rates.

Both young and old leaf tissue showed a significantly negative correlation between EXP and C(to), but this correlation was much stronger in old leaf tissue (**Figure 8 B, C**). In contrast, only in young leaf tissue, rates of NPS correlated positively with soluble carbohydrates and carboxylic acids while no significant correlation was observed in old leaf tissue. Further, sucrose and SPS activity were both negatively correlated with EXP only in old leaf tissue, while in young tissue no significant correlation was observed. This indicated a differential metabolic regulation of the total carbon balance in both types of tissue.

As these observations hinted towards a differential metabolic regulation of carbon uptake and release in young and old leaf tissue, a metabolic model was developed based on ordinary differential equations. The model described dynamics of sucrose turnover, carbon uptake, and export during the 8h light phase. To test whether the kinetic model could indicate a putative interaction of carbon metabolism in young and old leaf tissue, four flux scenarios were tested to account for potential sucrose exchange between young and old leaves. In the first scenario, both young and old metabolism was simulated independently describing the carbon balance equation as a non-specific export (*nsb*). In two further models, a directional flux from young to old (*y:o* model), or from old to young (*o:y* model) leaf tissue was implemented. Finally, in a third simulation scenario, a bidirectional exchange of sucrose between young and old tissue was assumed (*ambi* model). Model performance was assessed by the optimized cost function which represented least square errors between simulated and experimentally measured metabolite concentrations, i.e., the lower the cost function the better the fit of experimental data was.

Simulation of metabolism in Ct-1 and Oy-0 at 22 °C revealed best fits for *ambi* and *o:y* models (**Figure 9 A, C**). In Fei-0, simulations with lowest cost functions were realized by *ambi* and y:o models (**Figure 9 B**) while similar solution quality across all models was found for Rsch-4 at 22 °C (**Figure 9 D**).

**Figure 9.**
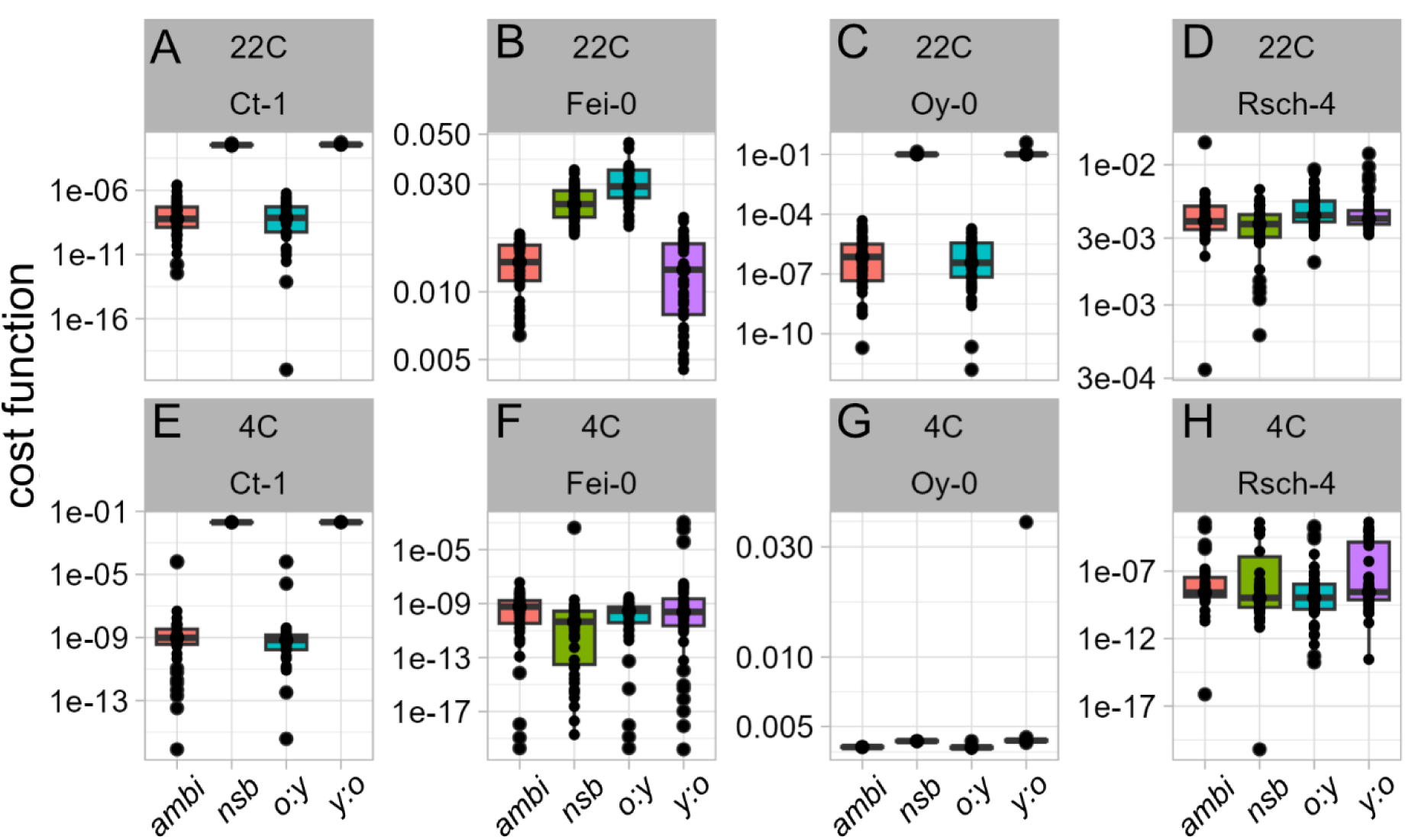
Cost functions of optimized kinetic models to simulate putative sucrose fluxes between young and old leaf tissue. **(A) – (D)** Decadic logarithm of least squared errors (cost functions) for accessions at 22 °C, **(E) – (H)** Decadic logarithm of least squared errors (cost functions) for accessions at LT/EL (4C). Each model was optimized 50x. red: ambidirectional sucrose transport (*ambi*); green; no sucrose transport between tissues (*nsb*); blue: sucrose transport from old to young tissue (*o:y*); purple: sucrose transport from young to old tissue (*y:o*). A graphical representation of the model structures is provided in Figure 1.

Similar cost function scenarios were observed at LT/EL, where *ambi* and *o:y* models fitted data best in Ct-1 and Oy-0 while model solution quality was indistinguishable in Rsch-4 (**Figure 9 E, G, H**). Hence, while all four tested model structures adequately captured the general diurnal dynamics of sucrose, glucose, and fructose concentrations in both young and old leaves under ambient and LT/HL conditions, the *ambi* and *o:y* assumptions yielded the most accurate representations of experimentally measured sucrose medians in old leaves (explicit solutions are shown in **Supplementary Figures S6 – S9**). Finally, however, the interpretation of model fitting in this study remains clearly limited by the low number of time points (eon, eod) which leaves considerable room for speculation about the dynamics during the day phase *in planta*.

## Discussion

A combination of low air temperature and elevated light intensities represents a frequently appearing challenging environment for plants, e.g., during sun rise on a cold morning. In such an environment, plants need to quickly and efficiently adjust photosynthesis and metabolism to prevent a harmful disbalance of light energy absorption, photosynthetic electron transport, enzymatic CO_2_ fixation and metabolism (Long et al., 1994). In the present study it was observed that *Arabidopsis* plants grown under short day conditions for eight to nine weeks show a gradient of F_v_/F_m_ under LT/EL across the leaf rosette. This finding was conserved across the four tested natural accessions which originated both from northern and southern European habitats. Hence, although freezing tolerance was not explicitly tested here, this indicates that such a gradient is a conserved trait which appears both in freezing sensitive and freezing tolerant accessions. Further analysis revealed that, already under ambient growth conditions, also net rates of CO_2_ uptake significantly differ across leaf rosettes where old leaves showed higher net assimilation rates than younger leaves. This might result from a lowered leaf respiration rate which has been shown earlier to decrease with leaf expansion, maybe due to a lowered number of mitochondria per unit cell volume (Armstrong et al., 2006). Such a developmental shift in mitochondrially located rather than in photosynthetic processes might also explain why F_v_/F_m_ did not differ between old and young tissue under ambient conditions. Interestingly, after exposure to LT/EL, both F_v_/F_m_ and rates of NPS were found to be higher in young than in old tissue. Although all measurements were conducted at 22 °C and, hence, do not represent the situation *in situ* (i.e., 4 °C), this suggested that LT/EL induced a photosynthetic and metabolic adjustment which changed the physiological role of old and young leaves when compared to ambient conditions. This was further reflected in the amounts of starch and sucrose which were found to be highest at eod in young leaf tissue across all accessions. While all accessions showed significant dynamics, i.e., accumulation, of sucrose during the light phase, starch accumulation was less conserved across accessions under LT/EL. Under these conditions, old leaf tissue of Ct-1 and Fei-0 was found to have reduced starch dynamics while in Oy-0 and Rsch-4 both tissue types showed a significant starch accumulation during the day. This might hint towards differential capacities of starch accumulation in tissue of putatively cold sensitive (Ct-1, Fei-0) and cold tolerant (Oy-0, Rsch-4) accessions. However, also regulation of starch degradation might be differentially regulated in the two considered tissues which has been shown to play a critical role in the development of cold and freezing tolerance (Kaplan and Guy, 2005; Sicher, 2011; Nagler et al., 2015). Starch metabolism has been discussed frequently in context of abiotic stress response and acclimation capacity, sometimes also with different dynamics across species (Thalmann and Santelia, 2017). While it remains speculation here, maybe developmental differences in leaf tissue contribute to these reported discrepancies which might also have different consequences for different stressors. As starch remobilization plays an important role under diverse conditions, such developmental effects might also become critical for (metabolic) engineering and breeding strategies.

In a recent study it was shown on the level of transcripts and proteins that, under changing growth light intensities and temperature regimes, old leaves support development and growth of younger leaves (Luklova et al., 2025). While the authors of this study applied low light intensities in combination with cold treatment, which contrasts the stress scenario of the present study, they showed that freezing tolerance was associated with lipid metabolism, chloroplast number and size. Independent of the underlying molecular and cellular mechanisms, also in the present study, which applied elevated light instead of low light, evidence was provided for a supportive role of mature leaves for growth and development of immature, or young, leaves. Correlation analysis of metabolites, enzyme activities and diurnal carbon balance revealed that net photosynthesis positively correlates with LT/EL-induced dynamics of sugars, carboxylic acids and enzyme activities of the central carbohydrate metabolism. This suggests that, in young leaves, an increased carbon assimilation rate is reflected by increasing carbon pools, e.g. carbohydrates and carboxylic acids, which might result in a stimulation of enzymatic carbohydrate interconversion. This contrasts findings of a previous study which showed that (maximum) enzyme activities only weakly, or not at all, correlated with metabolite levels across a large panel of *Arabidopsis* accessions (Sulpice et al., 2010). However, in this study, the authors did not apply a cold and/or light stress which shows that the observed correlations might heavily depend on the growth setup. Further, the present study provides compelling evidence for a differential correlation of enzymatic parameters, metabolites and carbon fluxes across one leaf rosette. For example, in old leaves, sucrose and the maximum activity of its biosynthesising enzyme SPS were found to negatively correlate with carbon export rates of the system, i.e., with the difference between input (NPS) and carbon turnover within the leaf. This was neither observed in a correlation analysis of young leaves nor in a correlation analysis which did not discriminate between old and young tissue (see **Figure 8**). This finding pointed towards a differential metabolic function of young and old leaf tissue when exposed to LT/EL. In general, young leaf tissue showed a strong increase of carbon uptake rates under LT/EL, but also old leaf tissue stayed (at least) constant compared to ambient conditions. Hence, metabolic changes, e.g., lower starch and sugar levels could not be (purely) explained by a lowered photosynthetic carbon assimilation. Also, amounts of carboxylic acids determined in old leaf tissue were similar to amounts of young tissue after exposure to LT/EL which supports the hypothesis that different correlations resulted rather from differential metabolic regulation than from net photosynthetic limitations. Finally, also a kinetic model with putative sucrose exchange between old and young leaf pointed towards a changing role of carbon supply under LT/EL: the best simulations of experimental data were gained in models which allowed for a sucrose exchange between both tissues, and preferentially from old to young leaves. While these findings point towards dynamic sink-source relationships between both tissues under stress (Roitsch, 1999; Lemoine et al., 2013), the question about the reason for such a dynamic remains elusive. Based on the finding that NPS rates in young leaf tissue can sufficiently supply and explain metabolite dynamics, even without sucrose exchange, it is hypothesised that the underlying plasticity of the metabolic system can equally well explain the observed metabolic phenotypes. Yet, as diurnal dynamics of most sugars under LT/EL were higher in young tissue than in old tissue, additional carbon import into the young metabolism could further improve the simulations.

Based on *in situ* stable isotope tracing it has previously been shown that sucrose utilization differs between *Arabidopsis* sink and source leaves (Dethloff et al., 2017). For example, citrate, fumarate and malate showed a higher label enrichment in sink than in source leaves. At least partially, this might also be reflected in the amounts of all three carboxylic acids quantified in the present study. Here, mature leaf tissue was observed to accumulate higher amounts of fumarate and malate under ambient conditions (see **Supplementary Figures S3, S4**). Interestingly, under LT/EL, especially malate amounts became similar across accessions and tissues, except for Fei-0 where older tissue still had higher levels than young tissue. Without discriminating mature and immature leaf tissue, such environmentally induced dynamics within leaf rosettes would be hidden behind the total variance of experimental data. Hence, to reduce ambiguity and to improve interpretability of leaf metabolomes, separated tissue analysis in large leaf rosettes might be beneficial. Finally, an obvious limitation of the present study is the low number of sampled time points across the light phase which reduces the discriminatory power between model simulation qualities, i.e., cost functions. However, together with the experimental data, the present study provides strong evidence for the need to separate *Arabidopsis* leaf rosettes in mature and immature parts if photosynthetic parameters, like F_v_/F_m_ or CO_2_ assimilation rates, indicate a gradient across leaf tissue. This might significantly improve interpretability and predictive power of quantitative models of metabolism which are developed to describe and predict plant growth, development and resilience in a changing environment.

## Acknowledgements

We thank the members of Plant Evolutionary Cell Biology at LMU München and the members of AG Weckwerth at University of Vienna for fruitful discussions. Further, we thank the Graduate School Life Science Munich (LSM) for support. This work was funded by Deutsche Forschungsgemeinschaft (DFG), NA1545-5/1.

## Data availability statement

Data of this study is available in the supplements.

## Author contributions

**VB**: Performed experiments, statistics, data evaluation and wrote the manuscript. **AK**: performed experiments and data evaluation. **MU**: performed experiments and data evaluation. **TN**: conceived the study, performed statistics and data evaluation, and wrote the manuscript.

## Supplementary Information

**Supplementary Figure S1.**
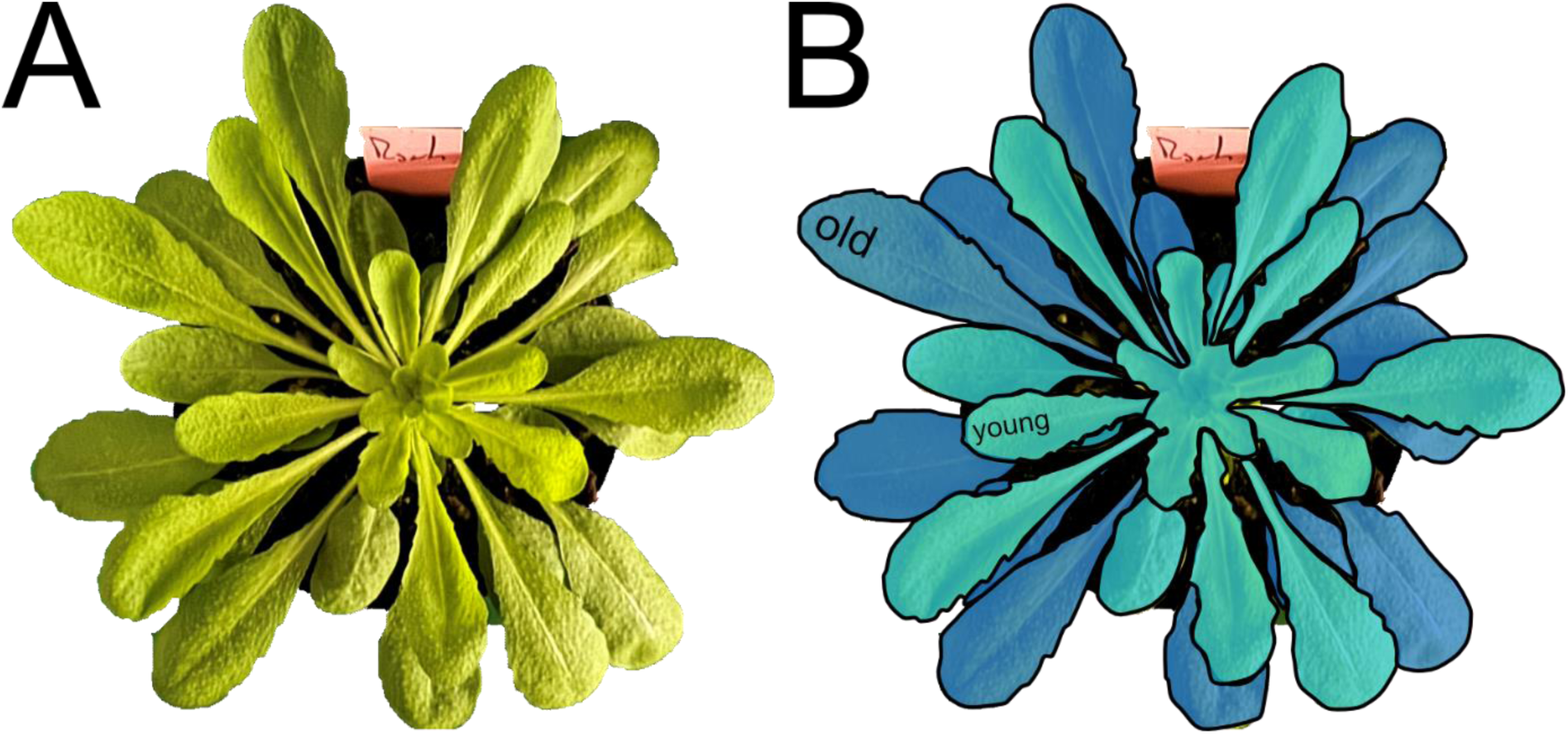
Classification of leaf tissue in a rosette of *Arabidopsis thaliana*. dark-blue: old/mature leaves; light-blue: young/immature leaves.

**Supplementary Figure S2.**
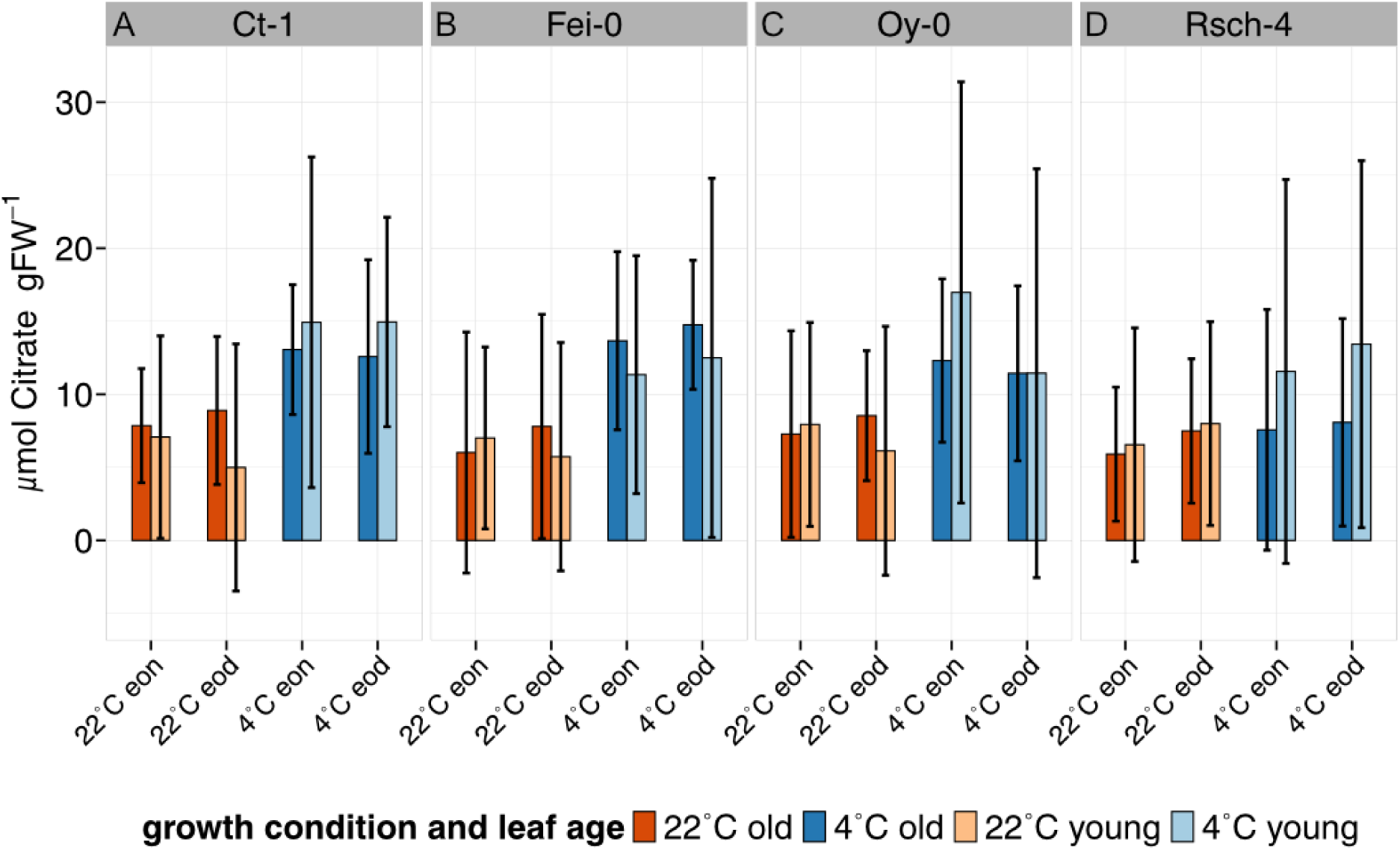
Dynamics of citrate amounts in young and old leaves of *A. thaliana.* **(A)** Citrate amounts in Ct-1, **(B)** Citrate amounts in Fei-0, **(C)** Citrate amounts in Oy-0, **(D)** Citrate amounts in Rsch-4. Bars represent means ± SD, n = 3. Orange: ambient temperature conditions, blue: 7d of LT/EL. Light shade: young leaves, dark shade: old leaves.

**Supplementary Figure S3.**
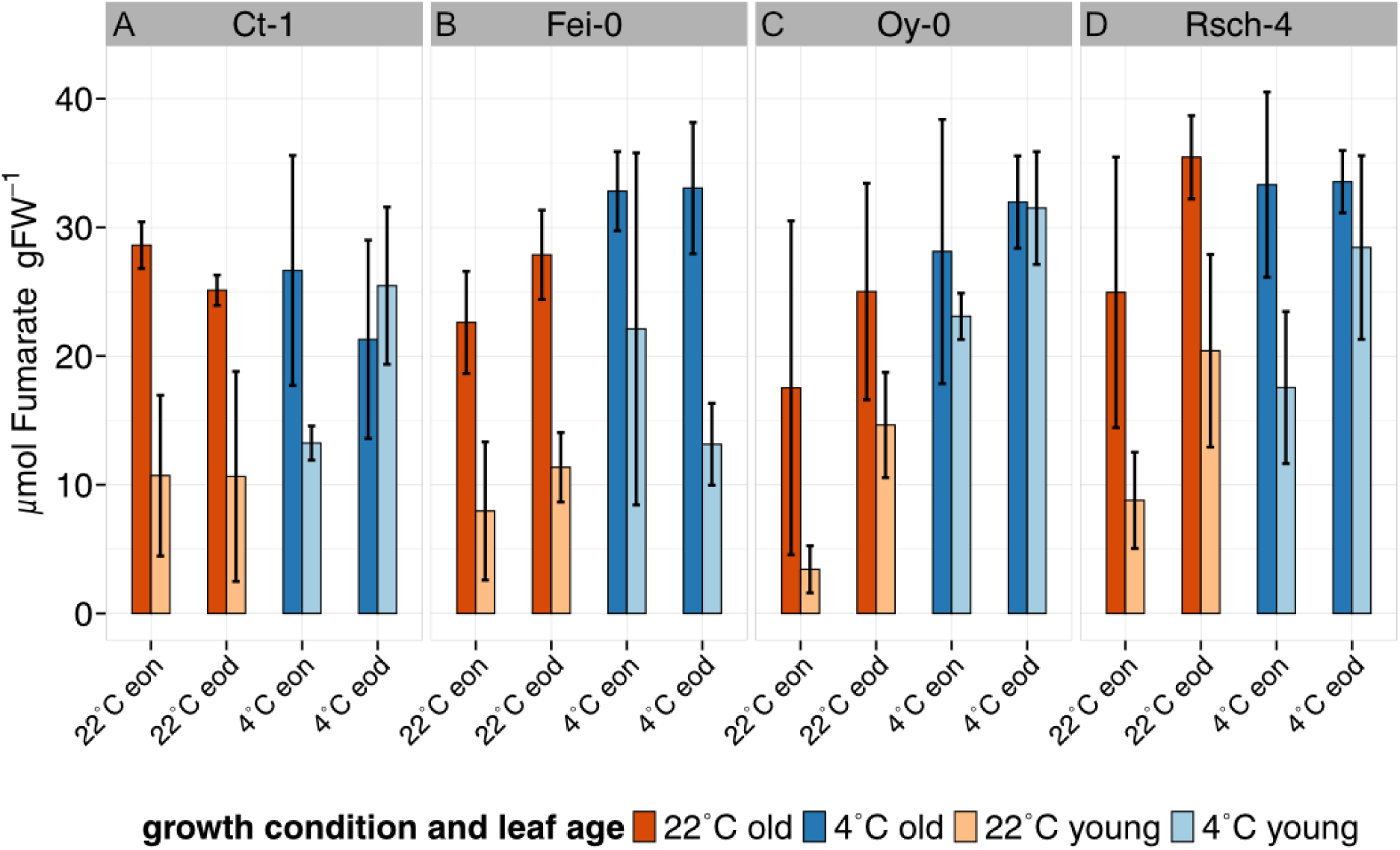
Dynamics of fumarate amounts in young and old leaves of *A. thaliana*. **A)** Fumarate amounts in Ct-1, **(B)** Fumarate amounts in Fei-0, **(C)** Fumarate amounts in Oy-0, **(D)** Fumarate amounts in Rsch-4. Bars represent means ± SD, n = 3. Orange: ambient temperature conditions, blue: 7d of LT/EL. Light shade: young leaves, dark shade: old leaves.

**Supplementary Figure S4.**
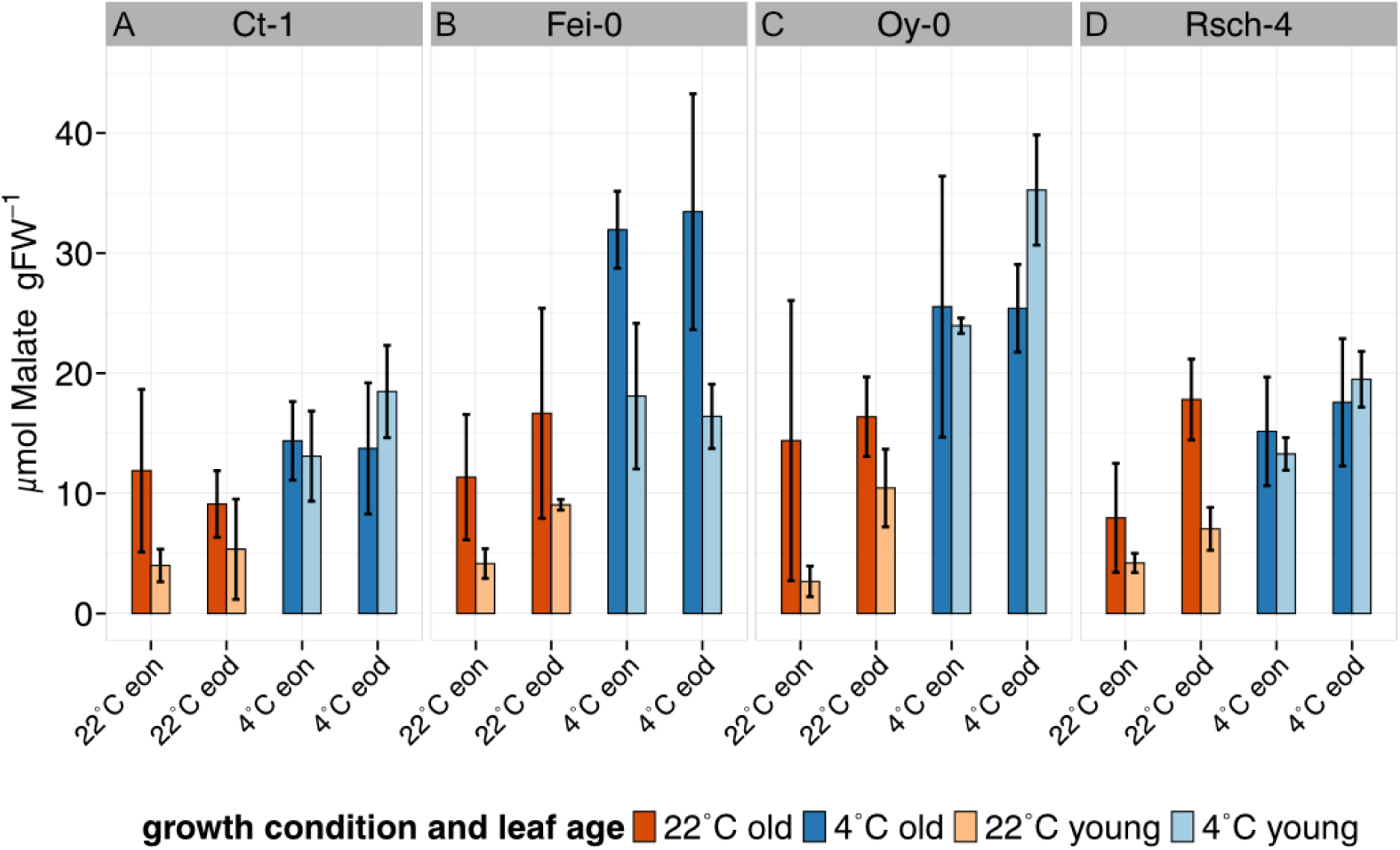
Dynamics of malate amounts in young and old leaves of *A. thaliana.* **(A)** Malate amounts in Ct-1, **(B)** Malate amounts in Fei-0, **(C)** Malate amounts in Oy-0, **(D)** Malate amounts in Rsch-4. Bars represent means ± SD, n = 3. Orange: ambient temperature conditions, blue: 7d of LT/EL. Light shade: young leaves, dark shade: old leaves.

**Supplementary Figure S5.**
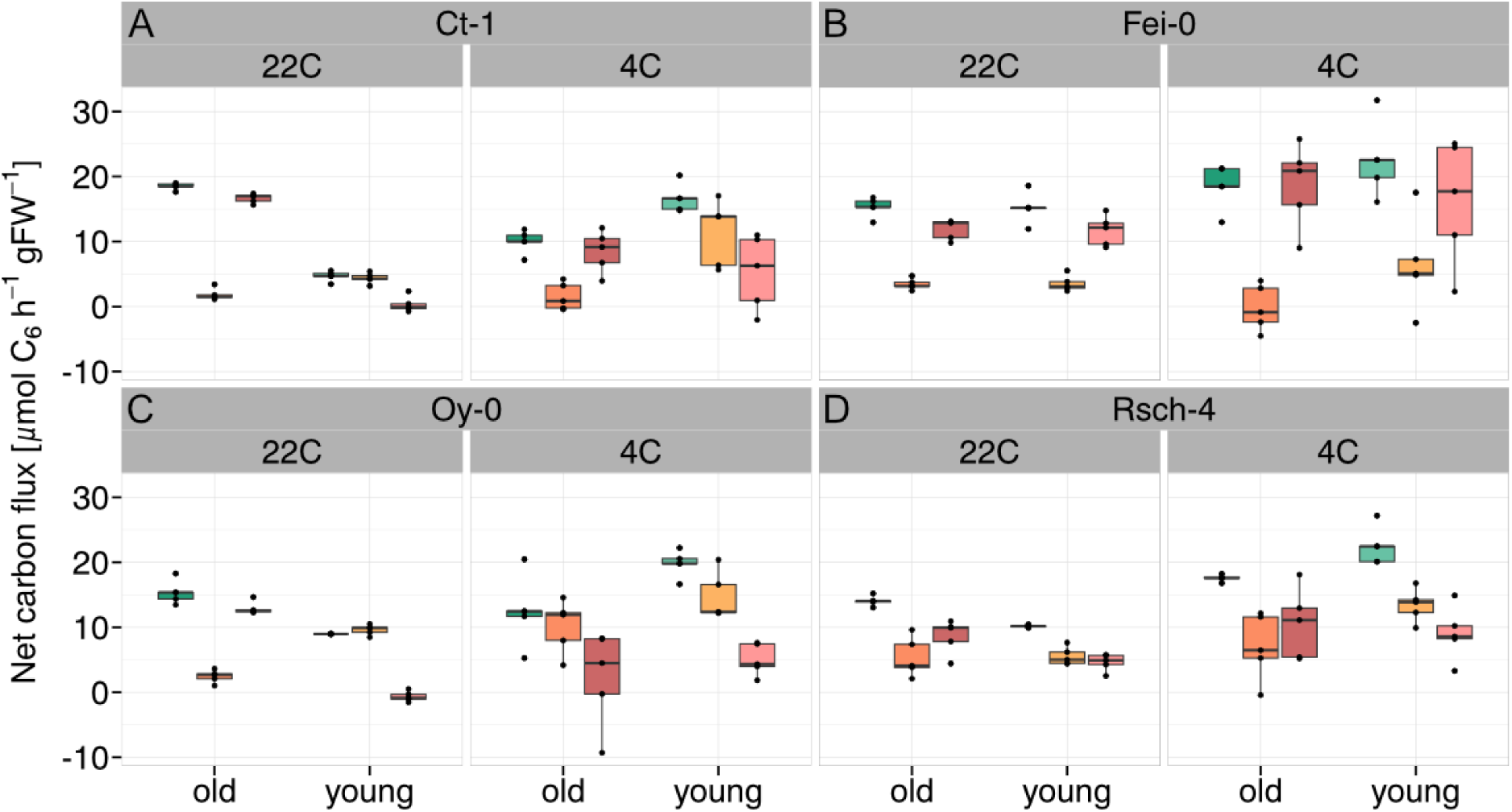
Balance between assimilated and metabolized carbon under ambient (22C) and LT/EL (4C) conditions. **(A)** Carbon fluxes in Ct-1, **(B)** Carbon fluxes in Fei-0, **(C)** Carbon fluxes in Oy-0, **(D)** Carbon fluxes in Rsch-4. Green: rate of assimilated carbon (NPS), orange: carbon turnover (C(to)) within leaves. pink: balanced output (NPS – C(to)).

**Supplementary Figure S6.**
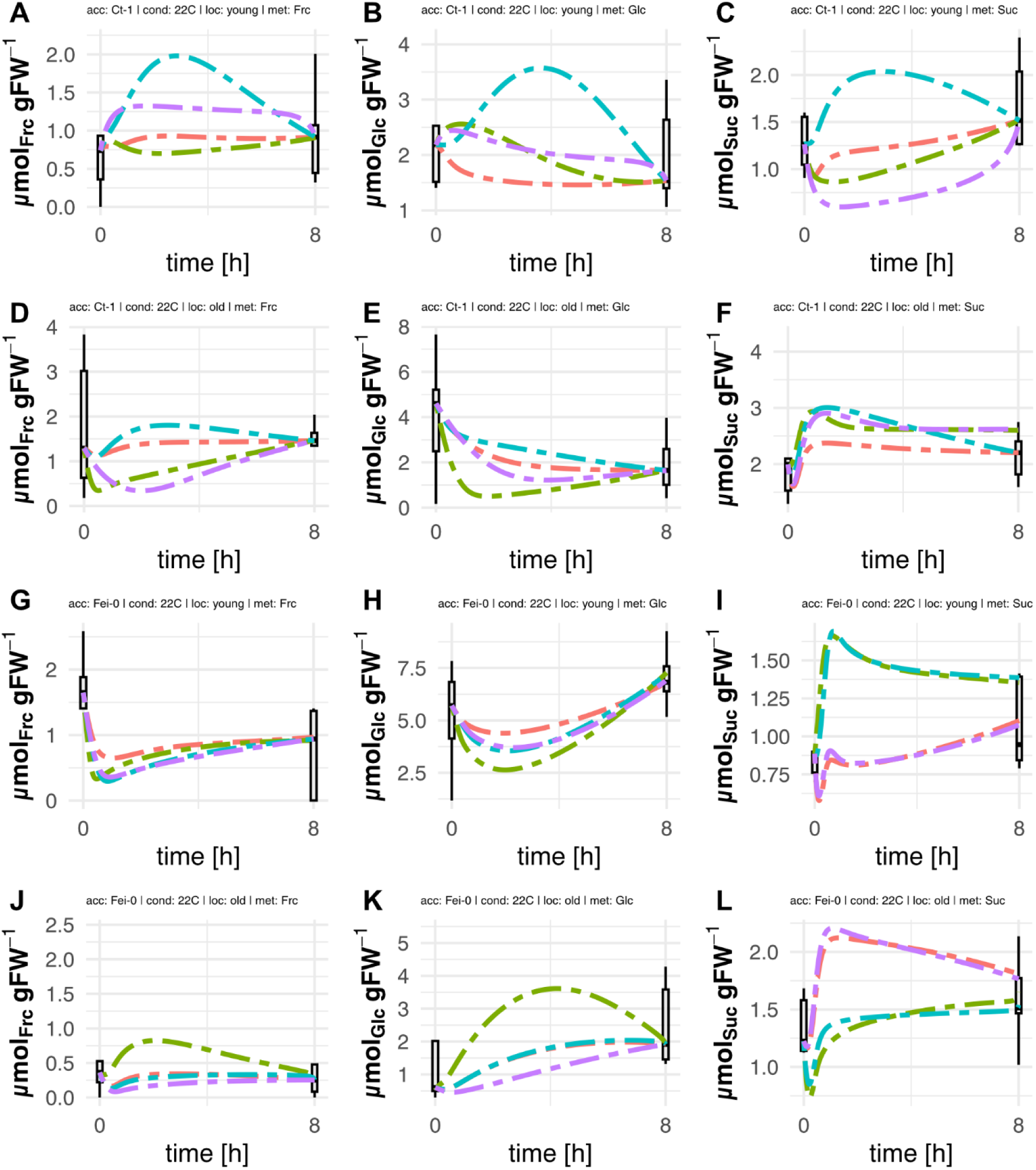
Simulated diurnal concentration changes in sucrose, glucose and fructose in old and young leaves of Ct-1 and Fei-0 at ambient conditions. **(A) – (C)** Simulations of sugar dynamics in Ct-1, 22 °C, young leaf tissue. **(D) – (F)** Simulations of sugar dynamics in Ct-1, 22°C, old leaf tissue. **(G) – (I)** Simulations of sugar dynamics in Fei-0, 22 °C, young leaf tissue. **(J) – (L)** Simulations of sugar dynamics in Fei-0, 22 °C, old leaf tissue. Boxes represent variance of experimental metabolite measurements (n = 5). Dashed lines indicate the best simulation results for metabolite concentrations (n = 50). Red: ambidirectional sucrose transport (*ambi*); green: no sucrose transport between leaves (*nsb*); blue: sucrose transport from old to young leaves (*o:y*); purple: sucrose transport from young to old leaves (*y:o*).

**Supplementary Figure S7.**
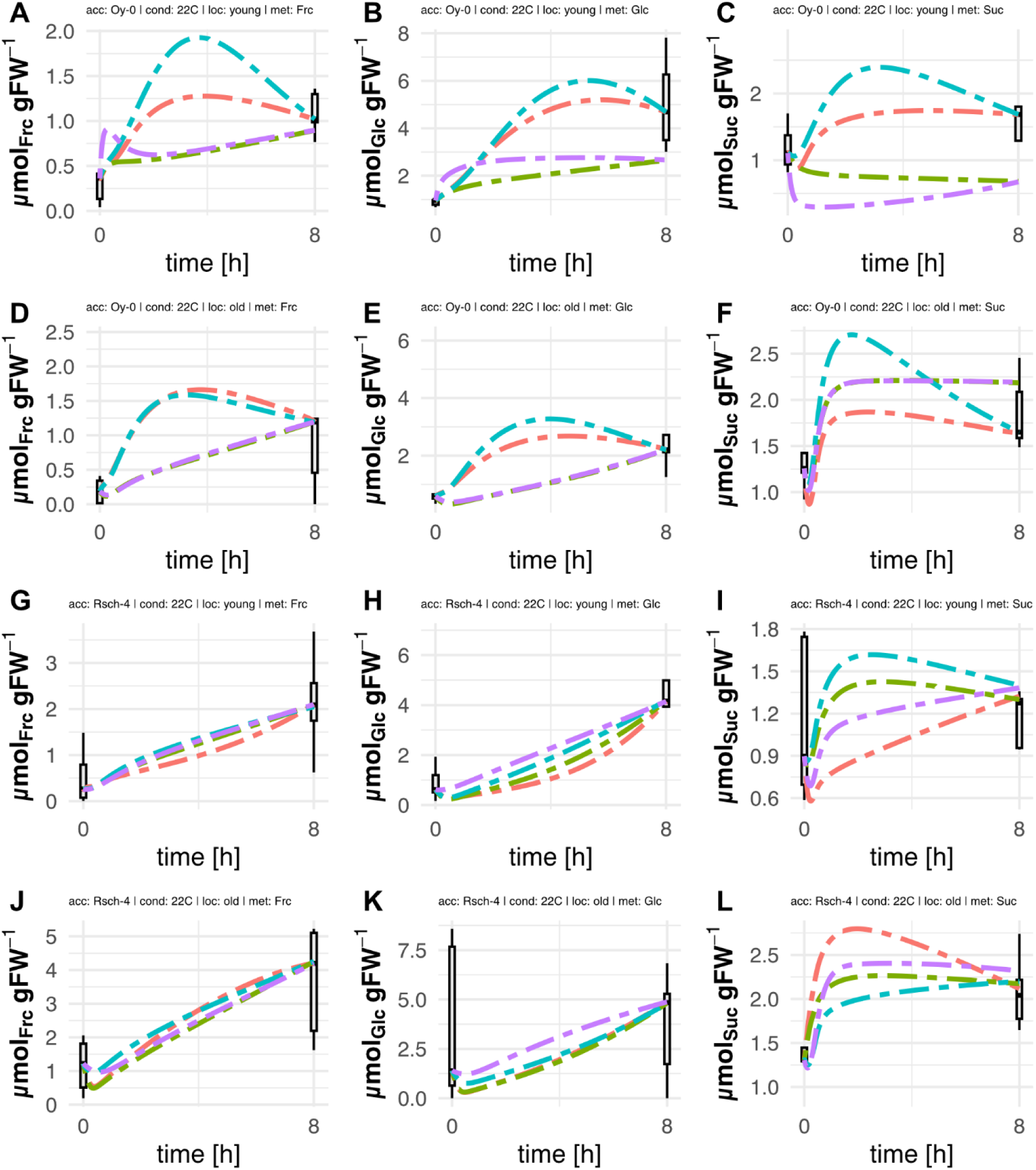
Simulated diurnal concentration changes in sucrose, glucose and fructose in old and young leaves of Oy-0 and Rsch-4 at ambient conditions. **(A) – (C)** Simulations of sugar dynamics in Oy-0, 22 °C, young leaf tissue. **(D) – (F)** Simulations of sugar dynamics in Oy-0, 22 °C, old leaf tissue. **(G) – (I)** Simulations of sugar dynamics in Rsch-4, 22 °C, young leaf tissue. **(J) –(L)** Simulations of sugar dynamics in Rsch-4, 22 °C, old leaf tissue. Boxes represent variance of experimental metabolite measurements (n = 5). Dashed lines indicate the best simulation results for metabolite concentrations (n = 50). Red: ambidirectional sucrose transport (*ambi*); green: no sucrose transport between leaves (*nsb*); blue: sucrose transport from old to young leaves (*o:y*); purple: sucrose transport from young to old leaves (*y:o*).

**Supplementary Figure S8.**
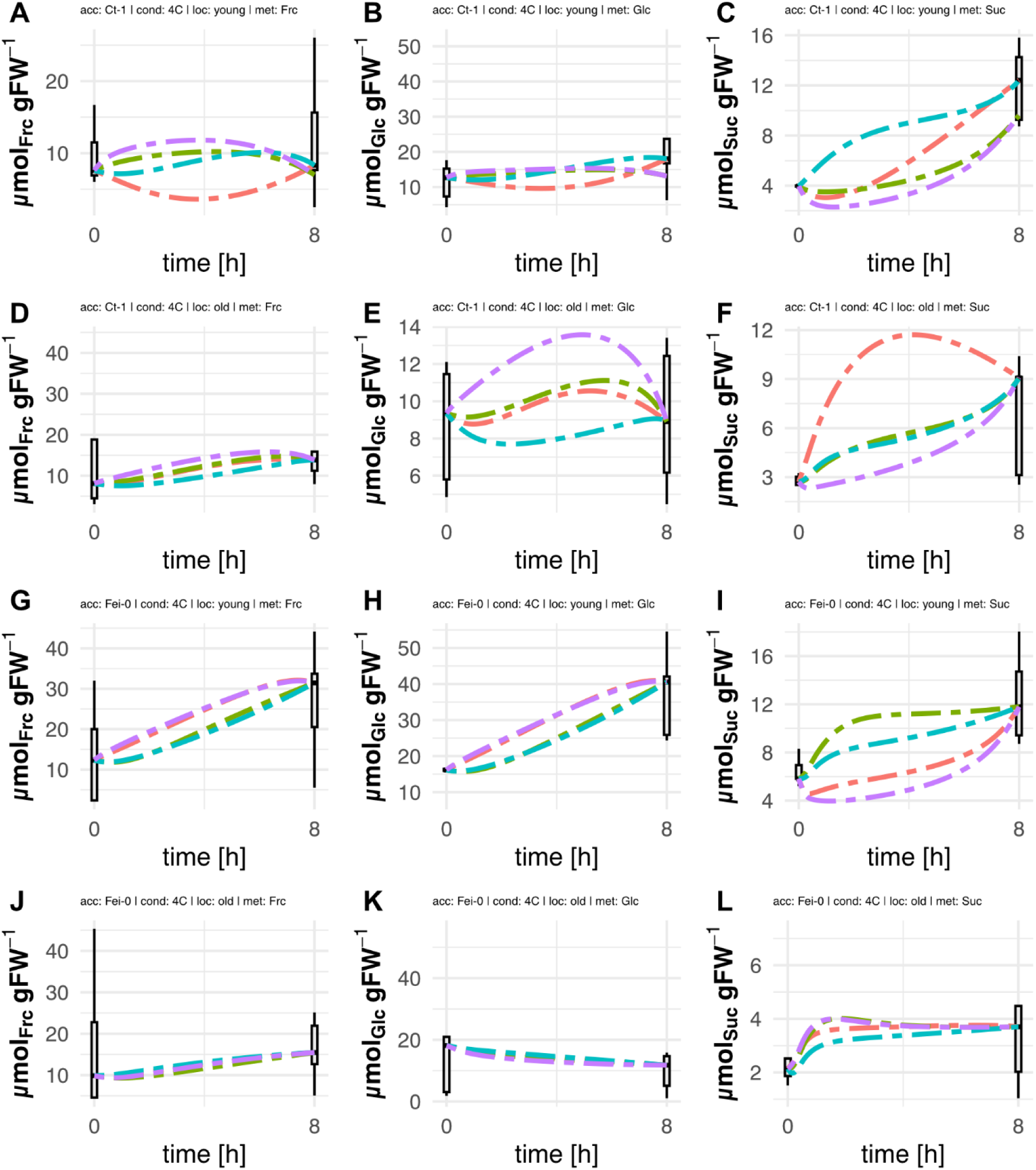
Simulated diurnal concentration changes in sucrose, glucose and fructose in old and young leaves of Ct-1 and Fei-0 at LT/EL. **(A) – (C)** Simulations of sugar dynamics in Ct-1, 4 °C, young leaf tissue. **(D) – (F)** Simulations of sugar dynamics in Ct-1, 4 °C, old leaf tissue. **(G) – (I)** Simulations of sugar dynamics in Fei-0, 4 °C, young leaf tissue. **(J) – (L)** Simulations of sugar dynamics in Fei-0, 4 °C, old leaf tissue. Boxes represent variance of experimental metabolite measurements (n = 5). Dashed lines indicate the best simulation results for metabolite concentrations (n = 50). Red: ambidirectional sucrose transport (*ambi*); green: no sucrose transport between leaves (*nsb*); blue: sucrose transport from old to young leaves (*o:y*); purple: sucrose transport from young to old leaves (*y:o*).

**Supplementary Figure S9.**
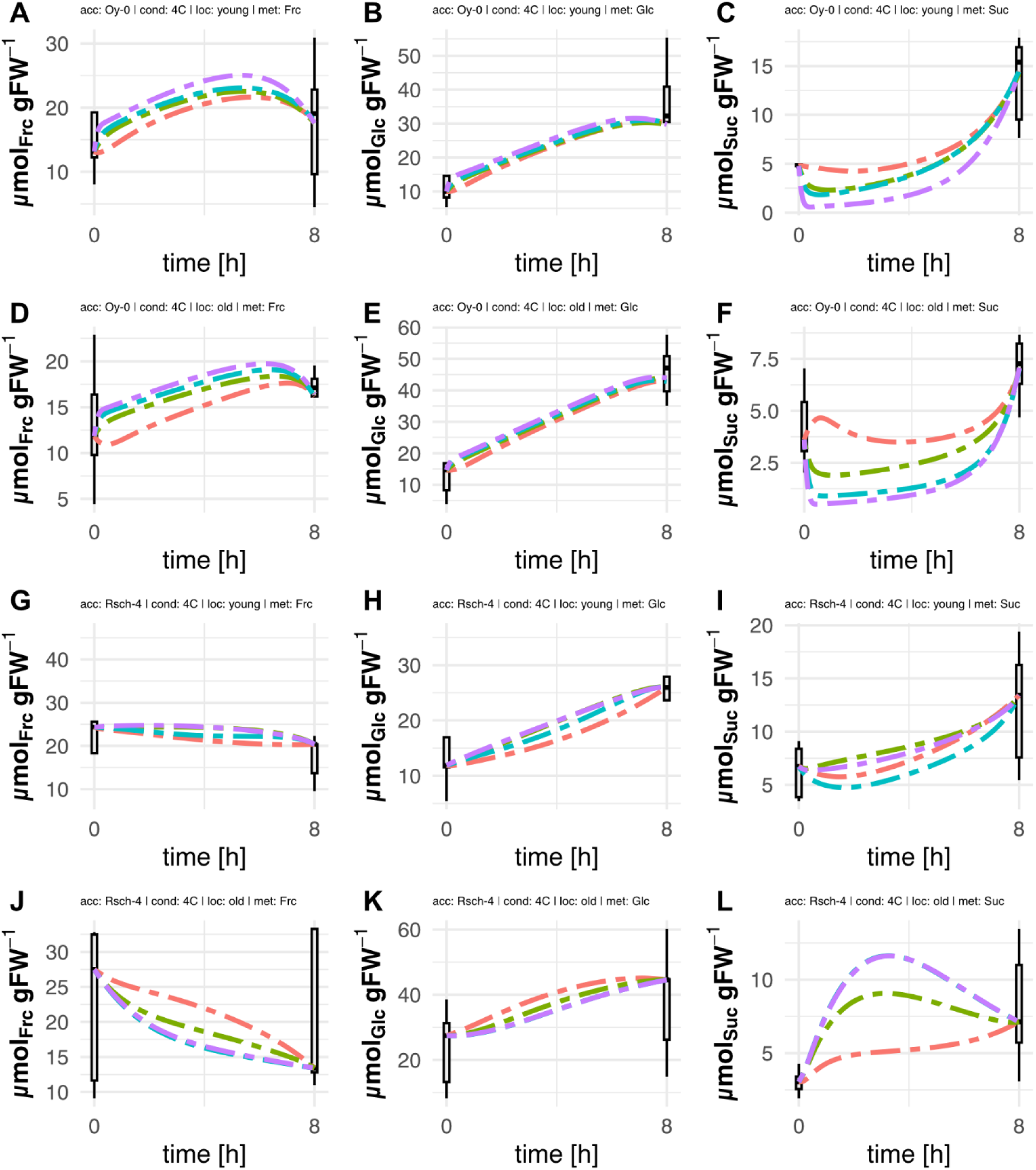
Simulated diurnal concentration changes in sucrose, glucose and fructose in old and young leaves of Oy-0 and Rsch-4 at LT/EL. **(A) – (C)** Simulations of sugar dynamics in Oy-0, 4 °C, young leaf tissue. **(D) – (F)** Simulations of sugar dynamics in Oy-0, 4 °C, old leaf tissue. **(G) – (I)** Simulations of sugar dynamics in Rsch-4, 4 °C, young leaf tissue. **(J) – (L)** Simulations of sugar dynamics in Rsch-4, 4 °C, old leaf tissue. Boxes represent variance of experimental metabolite measurements (n = 5). Dashed lines indicate the best simulation results for metabolite concentrations (n = 50). Red: ambidirectional sucrose transport (*ambi*); green: no sucrose transport between leaves (*nsb*); blue: sucrose transport from old to young leaves (*o:y*); purple: sucrose transport from young to old leaves (*y:o*).

**Supplementary Data F1. Model structure and parameters of all accessions and conditions.**

**Supplementary Data F2. Model parameters for best simulations**.

**Supplementary Data F3. Summary of all identified model parameters.**

