## Supplementary figures and images for "Acclimation of carbon metabolism to a changing environment across a leaf rosette of *Arabidopsis thaliana*"

### SF1_rosettes.tiff

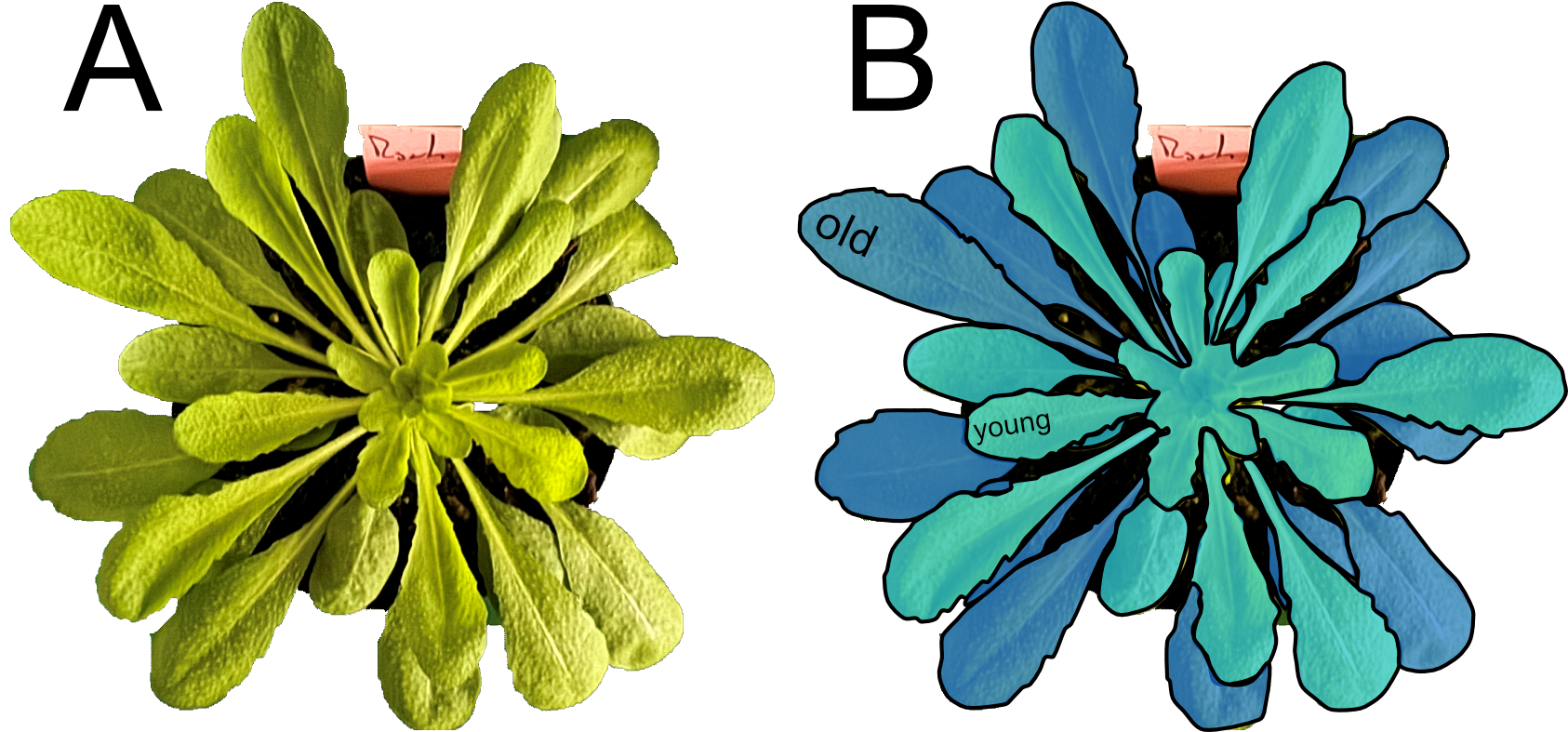

### SF2_Citrate_supp.tiff

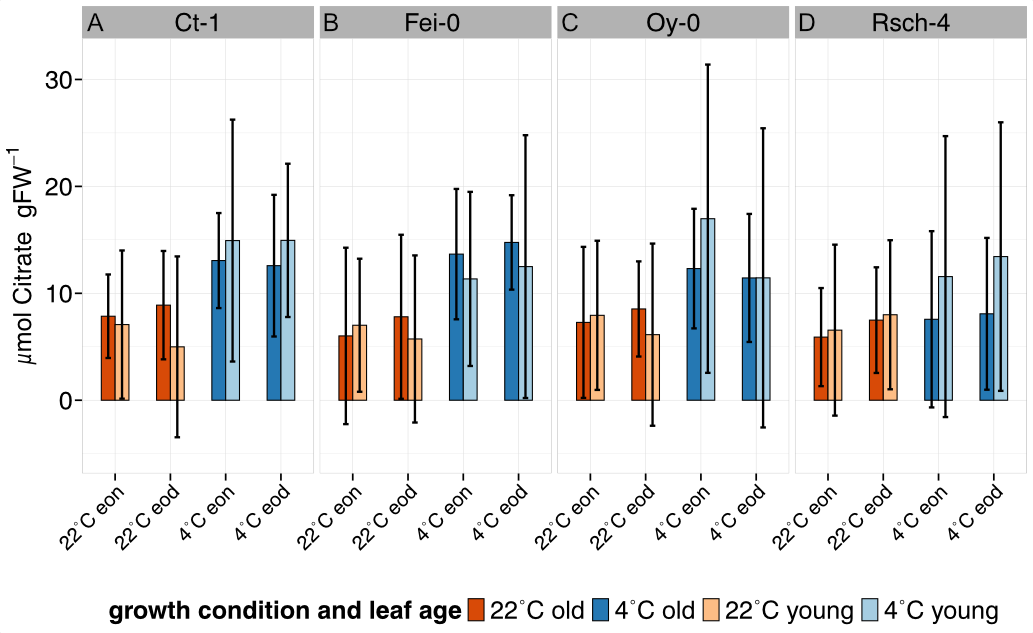

### SF3_Fumarate_supp.tiff

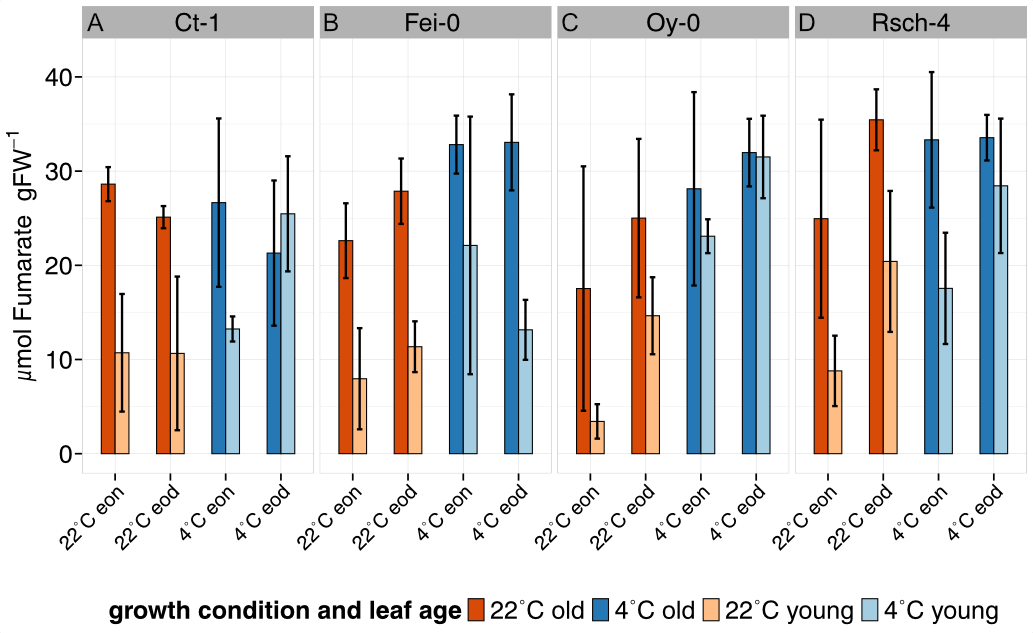

### SF4_Malate_supp.tiff

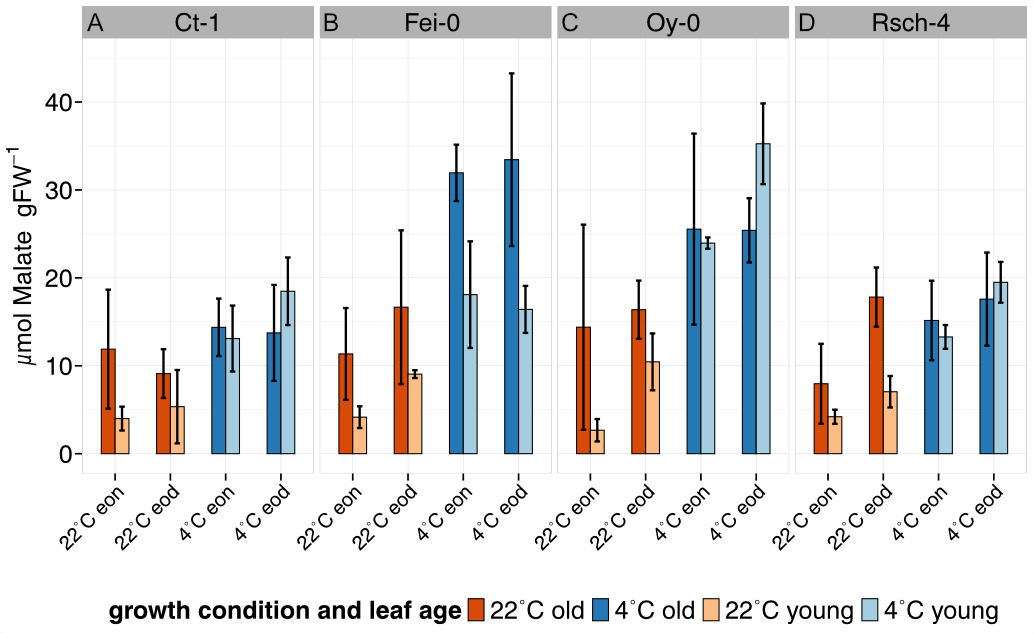

### SF5_balance model.tiff

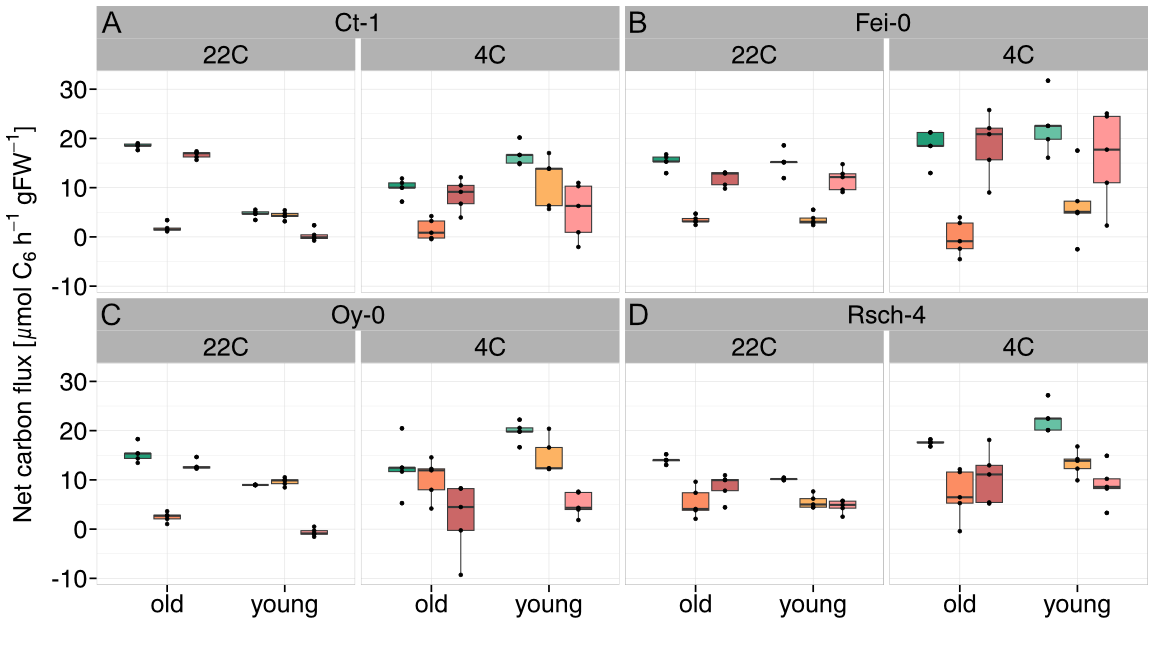

### SF6_simulations_ctfei_control.tiff

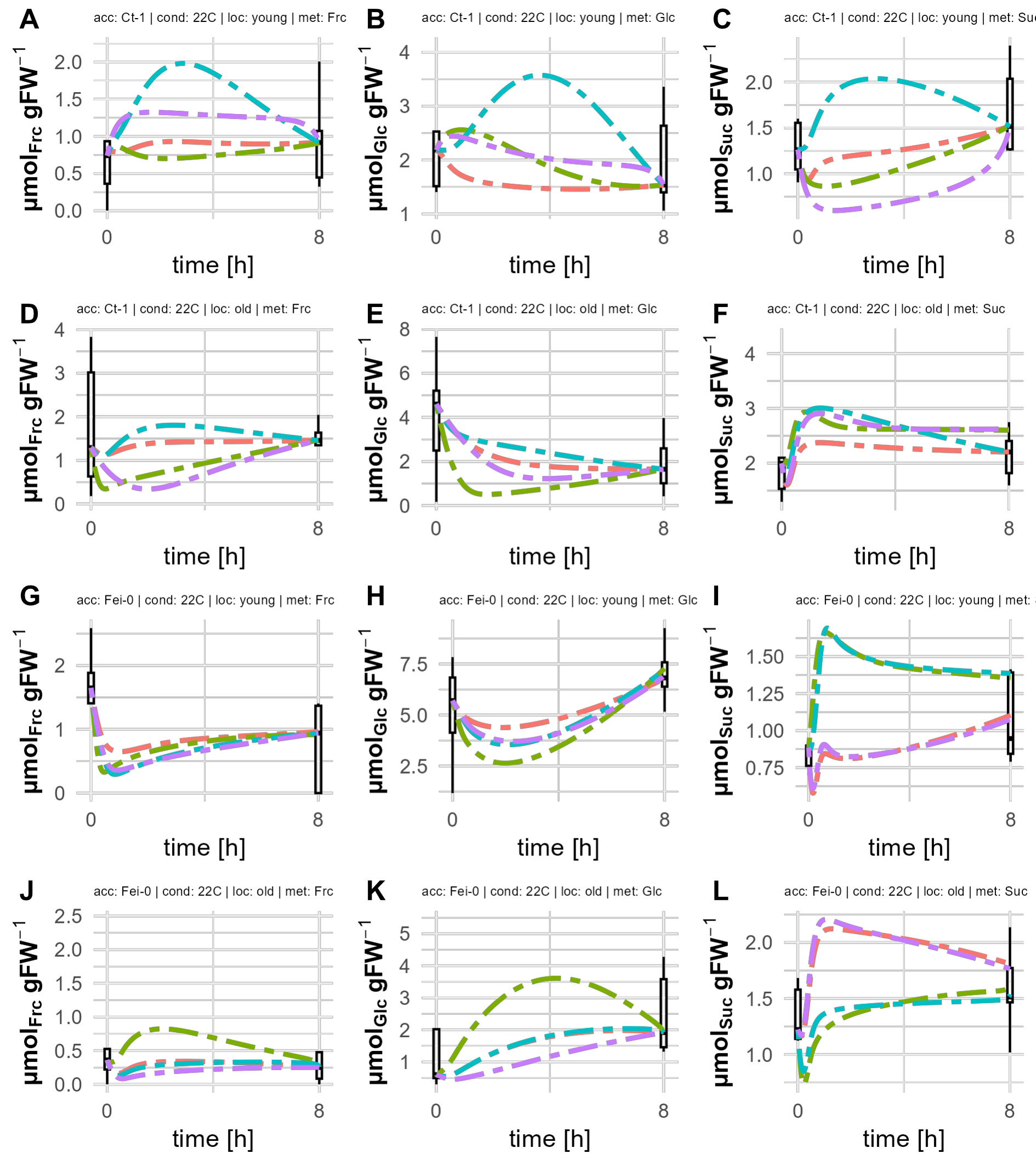

### SF7_simulations_oyrsch_control.tiff

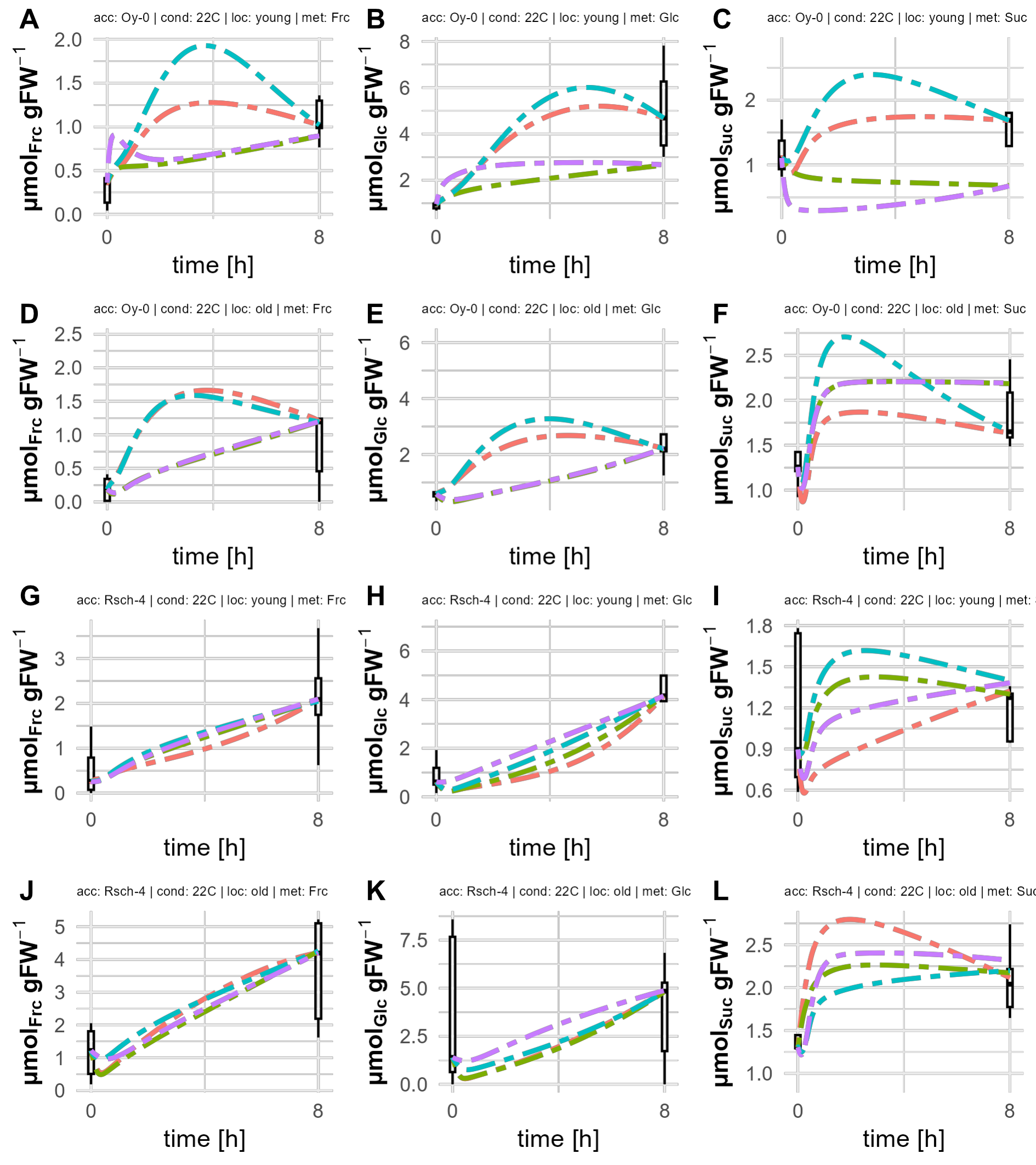

### SF8_simulations_ctfei_LT.tiff

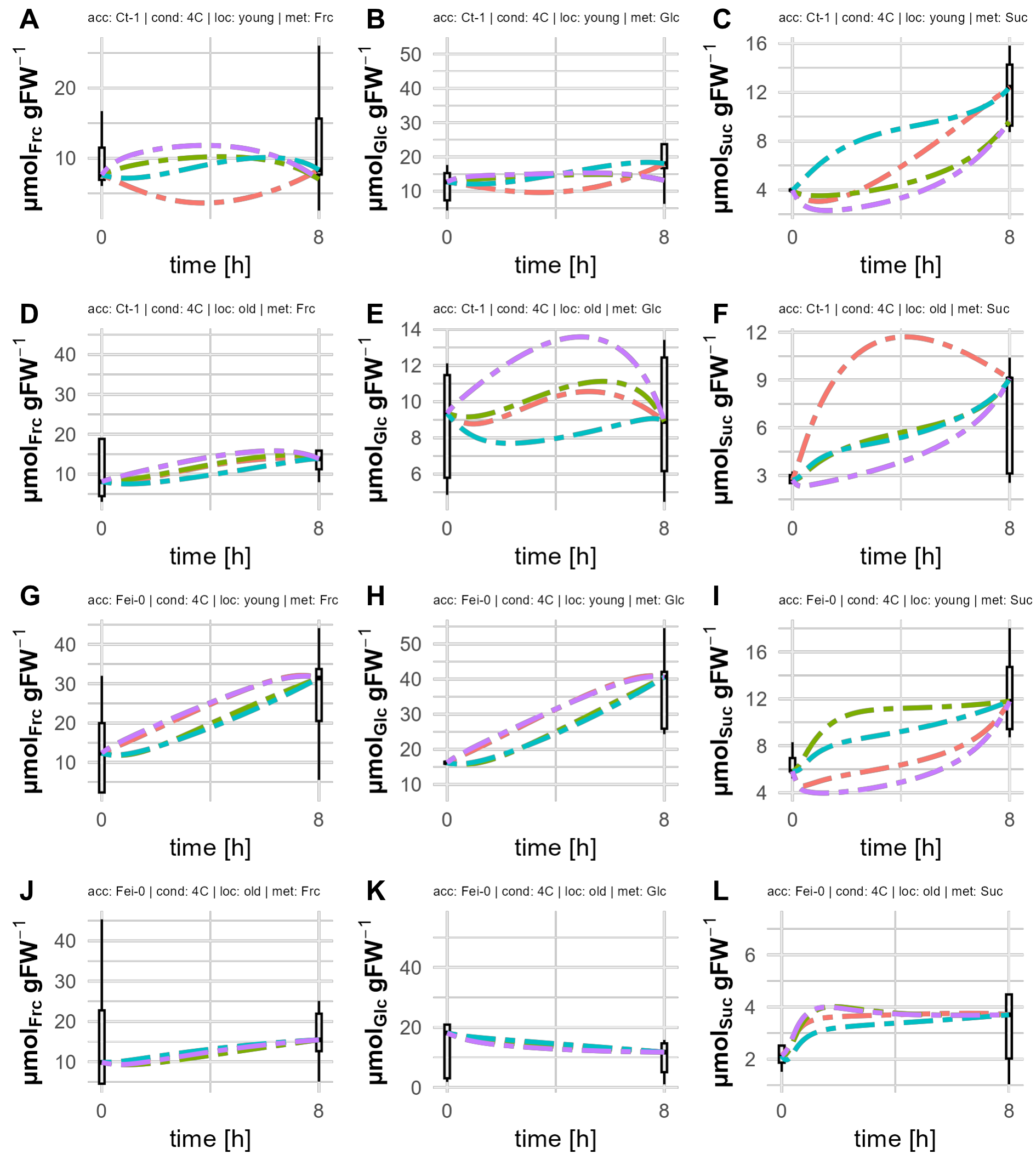

### SF9_simulations_oyrsch_LT.tiff

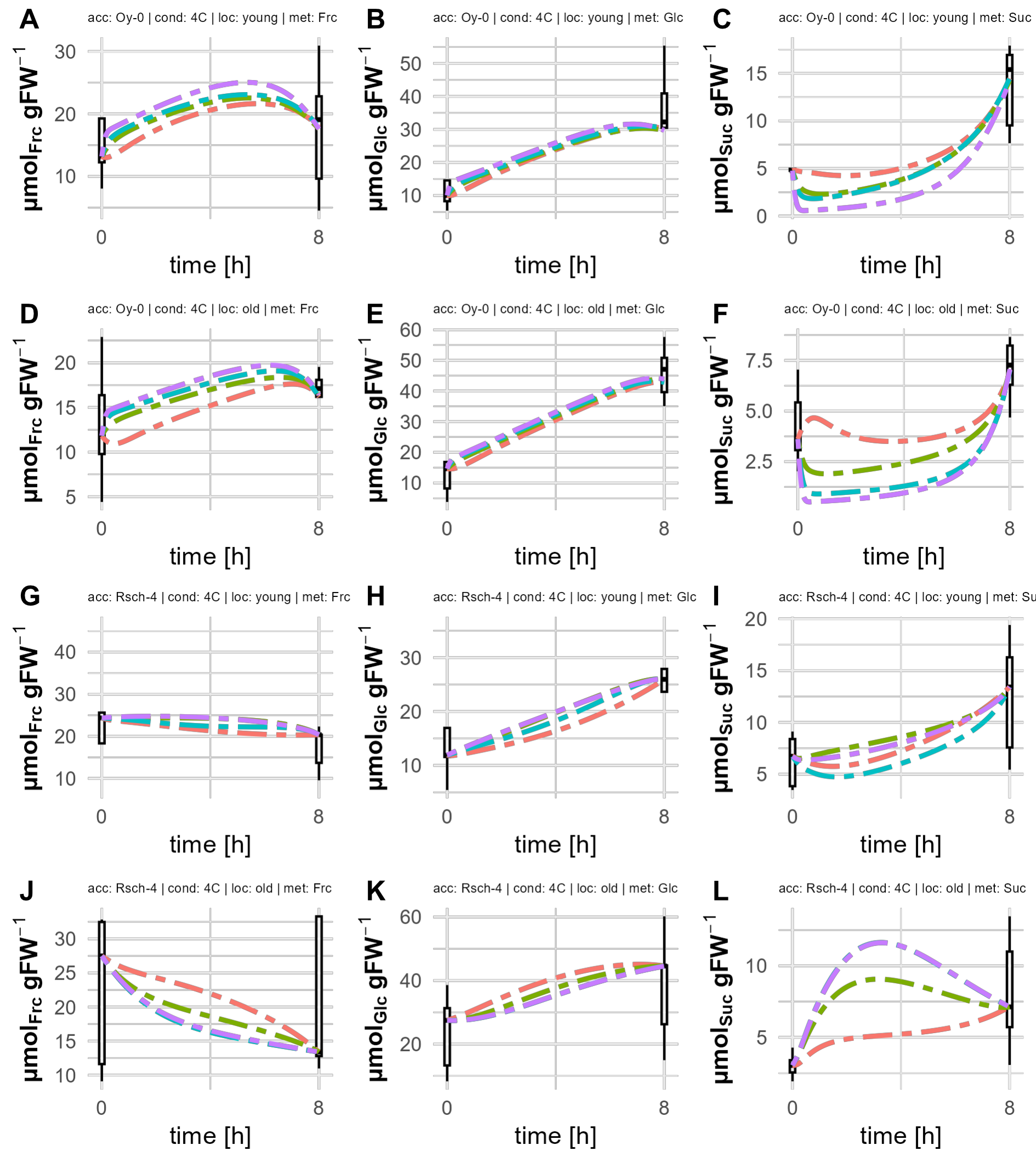
